# PH domain and leucine rich repeat phosphatase 1 (Phlpp1) suppresses parathyroid hormone receptor 1 (Pth1r) expression and signaling during bone growth

**DOI:** 10.1101/2020.12.06.413567

**Authors:** Samantha R. Weaver, Earnest L. Taylor, Elizabeth L. Zars, Katherine M. Arnold, Elizabeth W. Bradley, Jennifer J. Westendorf

**Author notes:** indicates equal contribution of authors (Samantha R. Weaver, Earnest L. Taylor). Corresponding author: Jennifer J. Westendorf, PhD, Mayo Clinic, 200 First Street SW, Rochester, MN 55905. **Disclosures** The authors declare that they have no disclosures.

## Abstract

Endochondral ossification is tightly controlled by a coordinated network of signaling cascades including parathyroid hormone (PTH). PH domain and leucine rich repeat phosphatase (Phlpp1) affects endochondral ossification by suppressing chondrocyte proliferation in the growth plate, longitudinal bone growth, and bone mineralization. As such, Phlpp1^−/−^ mice have shorter long bones, thicker growth plates, and proportionally larger growth plate proliferative zones. The goal of this study was to determine how Phlpp1 deficiency affects PTH signaling during bone growth. Transcriptomic analysis revealed greater Pth1r expression and H3K27ac enrichment at the Pth1r promoter in Phlpp1-deficient chondrocytes. PTH(1-34) enhanced and PTH(7-34) attenuated cell proliferation, cAMP signaling, CREB phosphorylation, and cell metabolic activity in Phlpp1-inhibited chondrocytes. To understand the role of Pth1r action in the endochondral phenotypes of Phlpp1-deficient mice, Phlpp1^−/−^ mice were injected with Pth1r ligand PTH(7-34) daily for the first four weeks of life. PTH(7-34) reversed the abnormal growth plate and long bone growth phenotypes of Phlpp1^−/−^ mice but did not rescue deficits in bone mineral density or trabecular number. These results demonstrate that elevated Pth1r expression and signaling contributes to increased proliferation in Phlpp1^−/−^ chondrocytes and shorter bones in Phlpp1-deficient mice. Our data reveal a novel molecular relationship between Phlpp1 and Pth1r in chondrocytes during growth plate development and longitudinal bone growth.

## INTRODUCTION

During development, long bones lengthen and harden through the process of endochondral ossification^(1)^. In the epiphyseal growth plate, chondrocytes proliferate and undergo hypertrophy to drive appositional bone lengthening. Vascularization of the early cartilaginous bone brings osteoblast and osteoclast precursors which ossify, model, and remodel bone. Most hypertrophic chondrocytes will undergo apoptosis, but some survive and contribute to the osteogenic pool in bone marrow^(2–4)^. These dynamic and complex processes are orchestrated by numerous extracellular factors that induce transcriptional events and intracellular signaling.

PH domain and leucine rich repeat phosphatases (Phlpp1 and Phlpp2) control cell proliferation and survival through posttranslational modification of several intracellular substrates^(5–7)^. Originally identified as terminators of Akt signaling^(8)^, further work showed Phlpp1/2 regulation of protein kinase C (PKC)^(9,10)^, ribosomal protein S6 kinase (S6K)^(11)^, and mitogen-activated protein kinase (Mapk/Erk)^(5)^. Additionally, Phlpp1/2 can translocate to the nucleus to modulate histone acetylation and phosphorylation^(12)^. Phlpp1/2 have been implicated as tumor suppressors^(9,13,14)^, but also regulate metabolic processes^(15)^, inflammation^(16)^, and tissue regeneration following injury^(17–24)^. Phlpp1 is overexpressed in human osteoarthritic tissue and Phlpp1 inactivation improves murine post-traumatic osteoarthritis^(17)^. Phlpp1 deletion accelerates chondrocyte maturation in vitro, preserves articular cartilage in vivo, and increases mobility in mice with post-traumatic osteoarthritis^(17,18)^. In the developing appendicular skeleton, Phlpp1 controls chondrocyte proliferation and bone lengthening^(5)^. Phlpp1-depleted mice have short long bones and low bone mass^(5)^ and Phlpp1-deficient chondrocytes proliferate more and express higher levels of growth factors such as Fgf18 and growth factor receptors, including Pth1r.

Pth1r is a G-protein-copuled receptor for parathyroid hormone (PTH) and parathyroid hormone-related protein (PTHrP). Dysregulation of signaling from Pth1r is at the root of several genetic diseases and is the target of bone anabolic therapies ^(25)^. Pth1r activates Gα subunits to regulate cyclic AMP production^(25)^, as well as Akt^(26)^, PKC^(27)^, and Erk1/2^(28)^ activation. Pth1r is expressed in various musculoskeletal cells, including mesenchymal stem cells^(29)^, osteocytes^(30)^, osteoblasts^(31,32)^, and chondrocytes^(33,34)^. In the growth plate, Pth1r activation in proliferating cells inhibits premature hypertrophy and thus is crucial for proper development prenatally and during early life^(1,35)^. PTH peptides (1-34 and 7-34) that bind Pth1r have been developed for clinical and research use. Both PTH(1-34) and PTH(7-34) cause Pth1r internalization with identical kinetics. However, only PTH(1-34) activates adenylyl cyclase and phospholipase C (PLC)^(36)^. When injected intermittently, PTH(1-34) increases bone mass and bone formation, and reduces fracture risk^(37)^. By contrast, continuous infusion of PTH(1-34) has catabolic effects on bone^(38)^. PTH(7-34) does not appear to have any effects on bone when administered alone^(39)^.

We previously showed that Pth1r mRNA levels were elevated in Phlpp1 depleted chondrocytes^(5)^. Here, we demonstrate that Phlpp1 regulates Pth1r transcription and signaling and show how these pathways cooperate to regulate endochondral ossification in mice. PTH(7-34) administration from birth through four weeks of age reversed the abnormal bone growth phenotype characteristic of Phlpp1^−/−^ mice. In vitro, PTH(7-34) reversed the increases in cell proliferation, CREB phosphorylation, metabolic activity, and cAMP signaling evident in Phlpp1^−/−^ chondrocytes. Taken together, our results demonstrate that Phlpp1 suppresses Pth1r expression and signaling in chondrocytes to regulate endochondral ossification.

## MATERIALS AND METHODS

### In vitro Phlpp inhibitor experiments

For experiments involving small-molecule Phlpp inhibitors (Phlpp*i*), cells were treated with 25μM NSC117079, 25μM NSC45586 (Glixx Laboratories), or vehicle (0.0005% DMSO) for the indicated time.

### In vitro PTH experiments

Primary WT or Phlpp1^−/−^ IMCs were plated and allowed to adhere overnight. Medium was replaced with DMEM containing either PTH(1-34), PTH(7-34) (Bachem), or vehicle (0.1% BSA in PBS) at the concentrations indicated.

### ATDC5 cell culture

ATDC5 cells were seeded at a density of 5 × 10^5^ cells/well in a six-well plate and incubated overnight to allow for cell adhesion in DMEM supplemented with 10% FBS and 100 units/mL penicillin, 100 μg/mL streptomycin, and 0.25 μg/mL Amphotericin B (1% antibiotic/antimycotic; Gibco 15240112). Phlpp*i* treatments were added as indicated. Each experiment included at least three technical replicates and was repeated at least three times. Results from a representative experiment are shown.

### Immature chondrocyte (IMC) cell culture

Primary immature chondrocytes (IMCs) were collected from 5-day-old WT or Phlpp1^−/−^ mice as previously described^(5,40)^. To evaluate mRNA and proteins, IMCs were isolated from cartilage with an overnight digestion in 0.5 mg/mL collagenase in serum-free culture medium, washed in PBS, and lysed as described below. When cells required treatment in culture, chondrocytes were plated in monolayer at a seeding density of 5 × 10^5^ cells/well in a six-well plate and cultured in DMEM supplemented with 2% FBS and 1% antibiotic/antimycotic. For micromass experiments, 10 μL drops of IMC suspensions (2 × 10^7^ cells/mL) were plated in DMEM. Three micromasses were plated in each well of a 6-well plate. After 1 hour, micromasses were covered with 2 mL DMEM + 2% FBS + 1% antibiotic/antimycotic and allowed to grow for 3 days, after which time the culture medium was changed to 2% FBS + 1% antibiotic/antimycotic + 1x Insulin/Transferrin/Selenium (ITS) + 0.05 mg/mL ascorbic acid + 10 mM β-glycerophosphate + respective treatment^(5,41)^. Littermates were pooled according to genotype for each experiment. Each experiment was repeated at least three times and represents at least n=3 biological replicates. Data from a representative experiment are shown.

### Transcriptional inhibition

Primary chondrocytes were plated and allowed to adhere overnight as described above. Actinomycin D (5μM, Gibco 11805017) was added for six hours and then replaced with medium containing Phlpp*i* or vehicle for 24 hours. Cells were collected for RNA or protein extraction.

### RNA isolation and real-time PCR

Total RNA was extracted from cell lines and primary chondrocytes using TRIzol (Invitrogen) and chloroform and 2μg was reverse transcribed using the iScript cDNA Synthesis Kit (Bio-Rad). Resulting cDNAs were used to assay gene expression via real-time PCR using the following gene-specific primers: Pth1r (5’-ACTTAGGCCGTTTCCTGTCC-3’, 5’-GAGGAGCTGACTCAGGTTGG-3’), Ywhaz (5’-GCCCTAAATGGTCTGTCACC-3’, 5’-GCTTTGGGTGTGACTTAGCC-3’). Fold changes in gene expression were calculated using the 2^−Δ ΔCt^ method relative to control after normalization of gene-specific Ct values to Ywhaz Ct values ^(42)^.

### Western blotting

Cell lysates were collected in a buffered SDS solution (0.1% glycerol, 0.01% SDS, 0.1M Tris, pH 6.8) on ice. Total protein concentrations were obtained using the Bio-Rad DC Assay (Bio-Rad). Proteins (20μg) were resolved by SDS-PAGE and transferred to a polyvinylidene difluoride membrane. Western blotting was performed with antibodies (1:1000 dilution) for Phlpp1 (Sigma 07-1341), Pth1r (Sigma SAB4502493), H3pS10 (Abcam ab5176), H3K9ac (Abcam ab10812), H3K27ac (Abcam ab4729), H3pS28 (Cell Signaling Technology 9713S), H3K9K14ac (Cell Signaling Technology 9677S), total H3 (Abcam ab1791), pCREB-S133 (Cell Signaling Technology 9198S), total CREB (Cell Signaling Technology 4820), and Actin (Sigma A4700) and corresponding secondary antibodies conjugated to horseradish peroxidase (HRP) (Cell Signaling Technology). Antibody binding was detected with the Supersignal West Femto Chemiluminescent Substrate (Pierce Technology, Rockford, IL).

### Live Cell Imaging

Primary chondrocytes (5 × 10^3^ cells/well) were allowed to adhere in monolayer in a 48-well plate and cultured overnight. Medium was replaced with DMEM containing either 10nM PTH(1-34), 10nM PTH(7-34), or vehicle (0.1% BSA in PBS). Cell confluency was detected in real-time with the IncuCyte S3 Live Cell Analysis System (Roche Applied Sciences, Indianapolis, IN), with four captures per well every hour for 48 hours.

### Cyclic AMP assay

WT and Phlpp1^−/−^ IMCs were allowed to attach to a 6-well plate overnight and then treated with 100nM PTH(1-34) or 100nM PTH(7-34) for 15 minutes. Cyclic AMP (cAMP) concentrations in IMC protein extracts (100 μg) were determined by ELISA (Cayman Chemical 581001).

### Chondrocyte metabolic activity assay

Primary chondrocytes (1×10^3^ cells/well) were cultured in monolayer in 96-well flat-well plates for 24 hours with 10nM PTH(1-34) or 10nM PTH(7-34). CellTiter 96® Aqueous One Solution Cell Proliferation Assays (Promega) were performed according to manufacturer specifications. Briefly, 20 μL of a reagent containing 3-(4,5-dimethylthiazol-2-yl)-5-(3-carboxymethoxy-phenyl-)-2-(4-sulfophenyl)-2H-tetrazolium (MTS) was added to each well for 24 h at 37□C. Spectrophotometer readings were taken at OD490.

### Micromass staining with Alcian Blue

Micromasses of WT or Phlpp1^−/−^ IMCs were plated as described above and incubated for three days in DMEM + 2% FBS. Medium was replaced with DMEM containing 1x ITS, 0.05 mg/mL ascorbic acid, and 10 mM β-glycerophosphate in the presence of 10nM PTH(1-34), 10nM PTH(7-34) for 9 days, with media changes every three days. Cells were washed with PBS, and 0.5% Alcian Blue (Sigma A5268) was applied for 2 hours.

### Histology and Immunohistochemistry / In Situ Hybridization

Right tibiae were decalcified for 14 days in 15% EDTA, dehydrated, and embedded in paraffin for sectioning. Sections (5 μm thick) were stained with Safranin O (Sigma S2255) and Fast Green (Sigma F7252)^(5)^ and chosen for analysis based on anatomical landmarks in the bone. Cross-sectional areas of the proliferative and hypertrophic zones in the proximal tibia growth plate were found in MatLab (version R2019b)^(43)^. A polygon was drawn around the respective zone and a binary image was formed where pixels inside the region of interest were set to one, and all other pixels were set to zero. The area of the binary image was calculated using the *bwarea* command and scaled to 3.45×10^−6^ mm^2^ per pixel. Immunohistochemistry was performed with antibodies diluted in 1% bovine serum albumin in tris-buffered saline directed to Pth1r (1:50 dilution, Sigma SAB4502493) or with a nonspecific IgG (control). Chromagens were detected with a polyvalent secondary HRP kit (Abcam, ab93697) and 3,3’-diaminobenzidine (DAB) (Sigma-Aldrich, D3939). In situ hybridization was performed using the RNAScope^®^ 2.5 HD Assay - Brown (ACD Biotechne). Briefly, slides were deparaffinized and epitope retrieval was achieved using a Custom Pretreatment Reagent provided by the manufacturer. Pth1r (ACD Biotechne 426191) and Col10a1 (ACD Biotechne 426181) riboprobes were hybridized to the tissue for two hours and detected using RNAScope^®^ 2.5 HD Assay detection reagents. DapB (ACD Biotechne 310043) was the negative control. Imaging was completed using a Zeiss LSM 900 Confocal Microscope.

### Chromatin Immunoprecipitation (ChIP) Sequencing and PCR

Primary chondrocytes were cultured in monolayer 10cm dishes for 24 hours. Cells were treated with 25μM NSC117079 or vehicle for 24 hours and then prepared for ChIP-Seq (2×10^7^ cells / sample). ChIP-seq was performed as previously described^(41)^, utilizing the anti-H3K27ac antibody (CST 8173BC (D5E4)). Libraries were prepared from 10ng DNA using the ThruPLEX DNA-seq Kit V2 (Rubicon Genomics, Ann Arbor, MI) and sequenced to 51 base pairs from both ends on an Illumina HiSeq 4000 instrument at the Mayo Clinic Medical Genome Facility Sequencing Core. ChIP-qPCR was performed as previously described utilizing 2 μg of anti-H3K27ac (Abcam ab4729) or YY1 (Active Motif 61780)^(41)^. Primers used for ChIP-qPCR were: Pth1r Promoter (5’-CCGCAGACTGACACGGAGAC-3’, 5’-CGACATTCATGGCAAGGCGG-3’), −1.47kb (5’-TGTGGAGTATCACACACTGCG-3’, 5’-TTGGGTAAAGCGGTCCCATT-3’). The −1.47kb primer is labeled to indicate the distance from the Pth1r promoter primer. The Pth1r promoter and upstream site were identified using Ensembl Genome Browser.

### PTH injections to mice

All mice were maintained in an accredited facility with 12-h light/dark cycle and supplied with food (PicoLab^®^ Rodent Diet 20 5058, LabDiet) *ad libitum* ^(44)^. All animal research was performed according to National Institute of Health and the Institute of Laboratory Animal Resources, National Research Council guidelines and the Mayo Clinic Institutional Animal Care and Use Committee approved all animal studies. Animals for these experiments were generated by crossing Phlpp1^+/−^ males and females. Beginning on the day after birth, entire litters of pups were injected daily with 100 μg/kg body weight/day PTH(7-34) or vehicle (0.1% BSA in PBS) through four weeks of age^(45)^. Animals were genotyped at three weeks of age^(5)^. Sample size was determined based on a power calculation that provided an 80% chance of detecting a significant difference (P<0.05). Final male group sizes were: WT+Vehicle (*n*=5), WT+PTH(7-34) (*n*=5), HET+Vehicle (*n*=9), HET+PTH(7-34) (*n*=6), Phlpp1^−/−^+Vehicle (*n*=4), and Phlpp1^−/−^+PTH(7-34) (*n*=6). Final female group sizes were: WT+Vehicle (*n*=7), WT+PTH(7-34) (*n*=6), HET+Vehicle (*n*=8), HET+PTH(7-34) (*n*=7), Phlpp1^−/−^+Vehicle (*n*=7), and Phlpp1^−/−^+PTH(7-34) (*n*=8). At four weeks old, mice were euthanized and limbs were collected, fixed overnight in 10% neutral buffered formalin, and stored in 70% ethanol until further analysis.

### X-ray Imaging

Radiographs of hind limbs were collected using a Faxitron X-ray imaging cabinet (Faxitron Bioptics, Tuscon, AZ). Limb length was measured on radiographs using ImageJ (1.52a) software.

### MicroCT

Micro-CT imaging of the femur and tibiae were performed using a SkyScan 1276 scanner (Bruker, Kontich, Belgium). Bones were fixed in 10% NBF before storage in 70% ethanol. Scans were performed at 55kV, 200 μA, 10 μm pixel resolution, 0.4□ rotation steps for 360□, 4 frames average imaging with a 0.25mm A1 filter. The acquired scans were reconstructed using the Skyscan NRecon software with beam hardening and post-alignment correction. Trabecular and cortical analyses of the femur were performed using Bruker CtAN software. The datasets were oriented in 3D to vertically align the longitudinal axis of each femur. As the bones were different lengths, a region of interest (ROI) for trabecular bone was defined as 5% the length of each bone, beginning 8% bone’s-length distance away from the distal growth plate. A gray-value threshold of 70 was applied to trabecular segmentations. Quantified outcomes were bone volume / total volume (BV/TV), trabecular thickness (TbTh), trabecular number (TbN), trabecular spacing (TbSp), and bone mineral density (BMD)^(46)^. For cortical bone analyses, the ROI was defined as 5% of total femur length beginning at the femoral midpoint. Quantified outcomes for cortical bone were tissue mineral density (TMD), cortical thickness (CTh), total tissue cross-sectional area (TtAr), cortical bone area (CtAr), and cortical area fraction (CtAr/TtAr)^(46)^. Semi-automatic segmentation of the proximal tibial growth plate was performed using 3D Slicer and a 3D region-growing method^(47)^ as previously described^(48)^. For one image slice, seeds were placed in the growth plate, bone, and empty space. This was repeated approximately 10 times in both the coronal and sagittal planes to define the segments in 3D. The “grow from seeds” feature was then applied to map the input image to user-specified segments. “Joint smoothing” was used on the resulting segmentation to preserve segment interfaces while removing noise. The growth plate segment was exported as a 3D patch object for processing in MatLab^(43)^. In MatLab, thickness measurements were taken by applying a query grid with 50 micron spacing, and sampling the z values of the patch object at the query points, resulting in approximately 3500 measurements across the growth plate. From these values, thickness distribution was plotted as a color map onto the patch object.

### Plasma and urine biochemical analyses

At four weeks of age, WT and Phlpp1^−/−^ mice were euthanized. Groups analyzed were: WT male (*n*=7), WT female (*n*=7), Phlpp1^−/−^ male (*n*=8), Phlpp1^−/−^ female (*n*=8). Whole blood was collected into EDTA-treated tubes (BD Biosciences 365974) and centrifuged for 10 minutes at 1,500 x g at 4°C to isolate plasma. Spot urine samples were collected immediately prior to euthanasia. Plasma intact PTH(1-84) was measured with an ELISA kit (Immutopics Inc., San Clemente, CA, USA). Urine creatinine and cAMP levels were measured via ELISA (Enzo Life Sciences, Farmingdale, NY, USA and Cayman Chemical, Ann Arbor, MI, USA). Urine was diluted 1:50 to fit within the standard curve of the creatinine kit and 1:300 for the cAMP kit.

### BrdU injections to mice and immunohistochemical analysis

Beginning on postnatal day 1 (P1), entire litters of pups from Phlpp1^+/−^ breeding pairs were injected with either 100 μg/kg body weight/day PTH(7-34) or vehicle (0.1% BSA in PBS) daily through P5. Males and females were combined because no sex differences were observed in earlier studies. The final group sizes were: WT+Vehicle (*n*=6), WT+PTH(7-34) (*n*=5), Phlpp1^−/−^+Vehicle (*n*=6), Phlpp1^−/−^+PTH(7-34) (*n*=8). On P5, pups were injected with 100 mL / 100 g body weight BrdU labeling reagent (Invitrogen 00-0103). Two hours later, pups were euthanized and hind limbs were collected, decalcified for 7 days in 15% EDTA, and processed for histology as described above. BrdU-positive cells were identified using the BrdU IHC kit (Millipore 2760). A representative area was chosen in the proximal tibia growth plate as described^(49)^, and the number of BrdU-positive cells per total cell number was calculated.

### Statistical analysis

Statistics were performed in Prism GraphPad (Version 8) using Student’s t-test, one-way ANOVA, or two-way ANOVA as appropriate with the necessary post-hoc tests for multiple comparisons. Data are depicted as individual points with SD bars (n=3) or boxplots showing the median, interquartile distance, and min/max values (n>3).

## RESULTS

### Phlpp1 inhibition increases Pth1r expression in chondrocytes

We previously reported that Pth1r mRNA levels are elevated in Phlpp1^−/−^ chondrocytes^(5)^. Given the important role of both Phlpp1 and Pth1r in endochondral bone formation, we sought to validate Pth1r expression changes in Phlpp1-inactivated cells. Phlpp inhibitors NSC117079 and NSC45586 increased Pth1r mRNA and protein expression in ATDC5 cells (**Figs 1A,B**; **Supp Fig 1A**) and IMCs (**Figs 1C,D; Supp Fig 1B**) within 24 hours. Elevated Pth1r expression was also confirmed by RT-PCR, immunoblotting, and immunohistochemistry in Phlpp1^−/−^ compared to WT mice (**Figs 1E-H; Supp Fig 1C**). In situ staining for Pth1r was more intense in the proximal tibial growth plate of Phlpp1^−/−^ mice than WT mice at both the RNA and protein levels (**Fig 1G,H; Supp Fig 2A**). Thus, Phlpp1 inhibition rapidly elevates Pth1r expression in vitro and in vivo.

**Figure 1.**
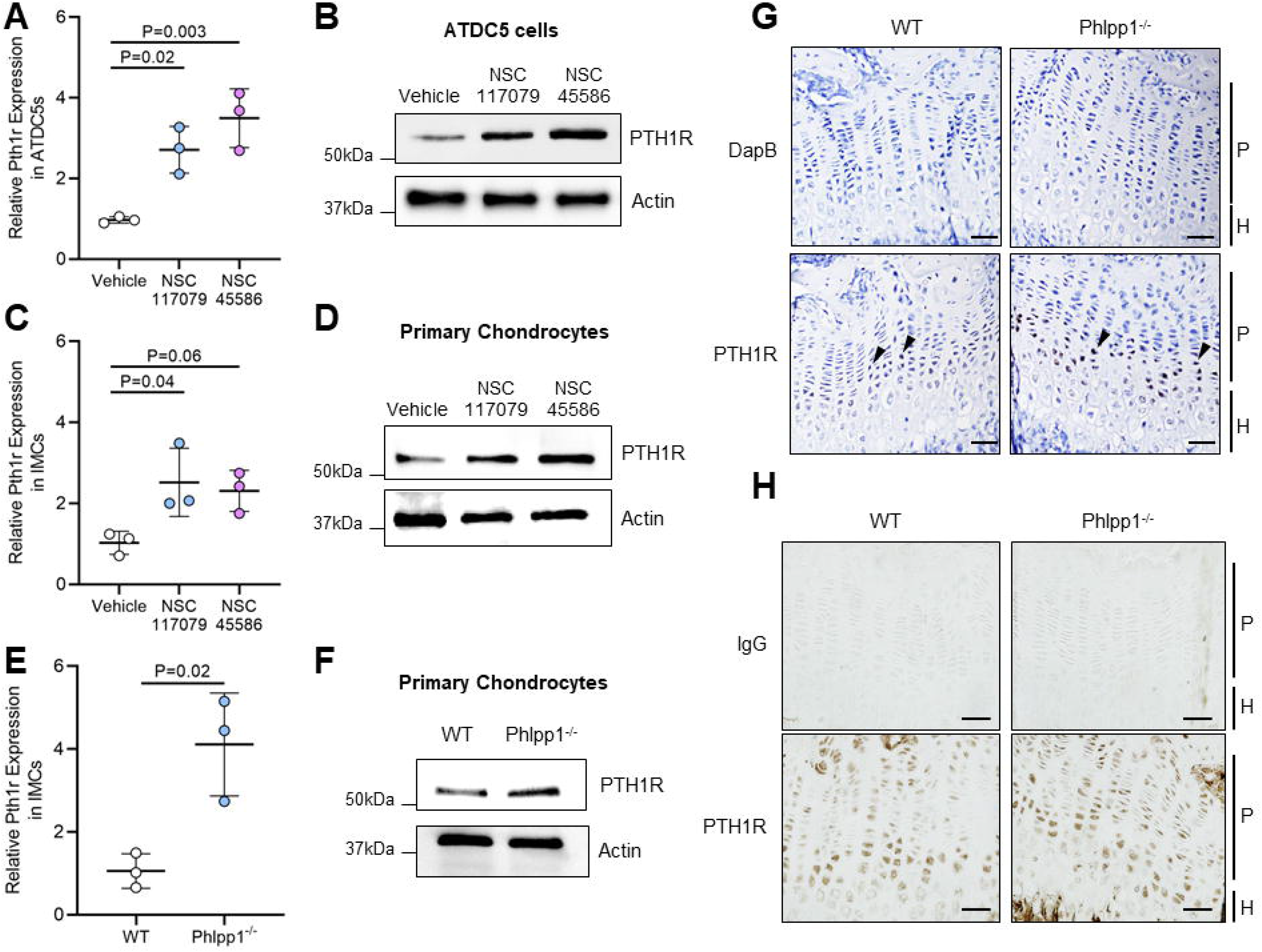
Phlpp inhibition increases Pth1r expression in chondrocytes. (A-D) Pth1r mRNA expression and protein levels were measured in ATDC5 cells (A,B) and primary chondrocytes (C,D) cultured in the presence of Phlpp inhibitors for 24 hours. Pth1r (E) mRNA expression and (F) protein levels were measured in Phlpp1^−/−^ primary chondrocytes compared to WT. (G,H) The proximal tibial growth plates of 4-week-old male WT and Phlpp1^−/−^ mice are shown. (G) In situ hybridization was performed with a riboprobe targeted to Pth1r. Scale bar = 50 μm. (H) Immunohistochemistry was performed with an antibody targeted to Pth1r. Scale bar = 20 μm. Vertical black lines indicate either P = proliferative zone or H = hypertrophic zone. Statistically significant differences were determined with one-way ANOVA with Tukey’s post-hoc test (A, C) or Student’s t test (E).

### Phlpp1 regulates Pth1r through transcription

We next identified mechanisms by which Phlpp1 inactivation increases Pth1r levels. Phlpp1 is known to modulate histone 3 (H3) acetylation and phosphorylation ^(12,50)^. Phlpp inhibitors NSC117079 and NSC45586 increased phosphorylation (p) of H3S10 (H3pS10) as well as acetylation (ac) of H3K9 and H3K27 in primary IMCs, but not H3pS28 or H3K9K14ac (**Fig 2A; Supp Fig 1D-H**). Similar results were observed in Phlpp1^−/−^ IMCs (**Fig 2A; Supp Fig 1I-M**). ChIP-seq revealed several H3K27ac-enriched regions in control and NSC117079-treated cells. The abundance of H3K27ac was greater within the Pth1r promoter in IMCs treated with the Phlpp inhibitor NSC117079 (**Fig 2B**). When compared to publicly available datasets, H3K27ac peaks are similar to those found on embryonic limbs (**Supp Fig 3**)^(51)^. ChIP-PCR of chromatin from IMCs treated with NSC117079 and NSC45586 for 24 hours verified robust enrichment of H3K27ac in the Pth1r promoter of IMCs (**Fig 2C**). Phlpp1^−/−^ IMCs also had greater basal enrichment of H3K27ac in the Pth1r promoter compared to WT IMCs (**Fig 2D**). The RNA polymerase inhibitor, actinomycin D, prevented the increase in Pth1r mRNA and protein levels, indicating transcriptional regulation (**Fig 2F; Supp Fig 1N**). A search of two databases identified binding sites for over 70 transcription factors in the region of the Pth1r promoter that was hyperacetylated after Phlpp1 inhibition (**Supp Table 1**). The transcription factor Ying Yang 1 (YY1) was chosen for further evaluation because it was present in both databases, had multiple potential binding sites in the H3K27ac-enriched peak of the Pth1r promoter, and was detectable in primary chondrocytes via qPCR (data not shown). Primary chondrocytes treated with the Phlpp inhibitor NSC117079 for 30 minutes had increased binding of YY1 in the Pth1r promoter (**Fig 2F**). Together, these data demonstrate that Phlpp inactivation increases transcription of Pth1r.

**Figure 2.**
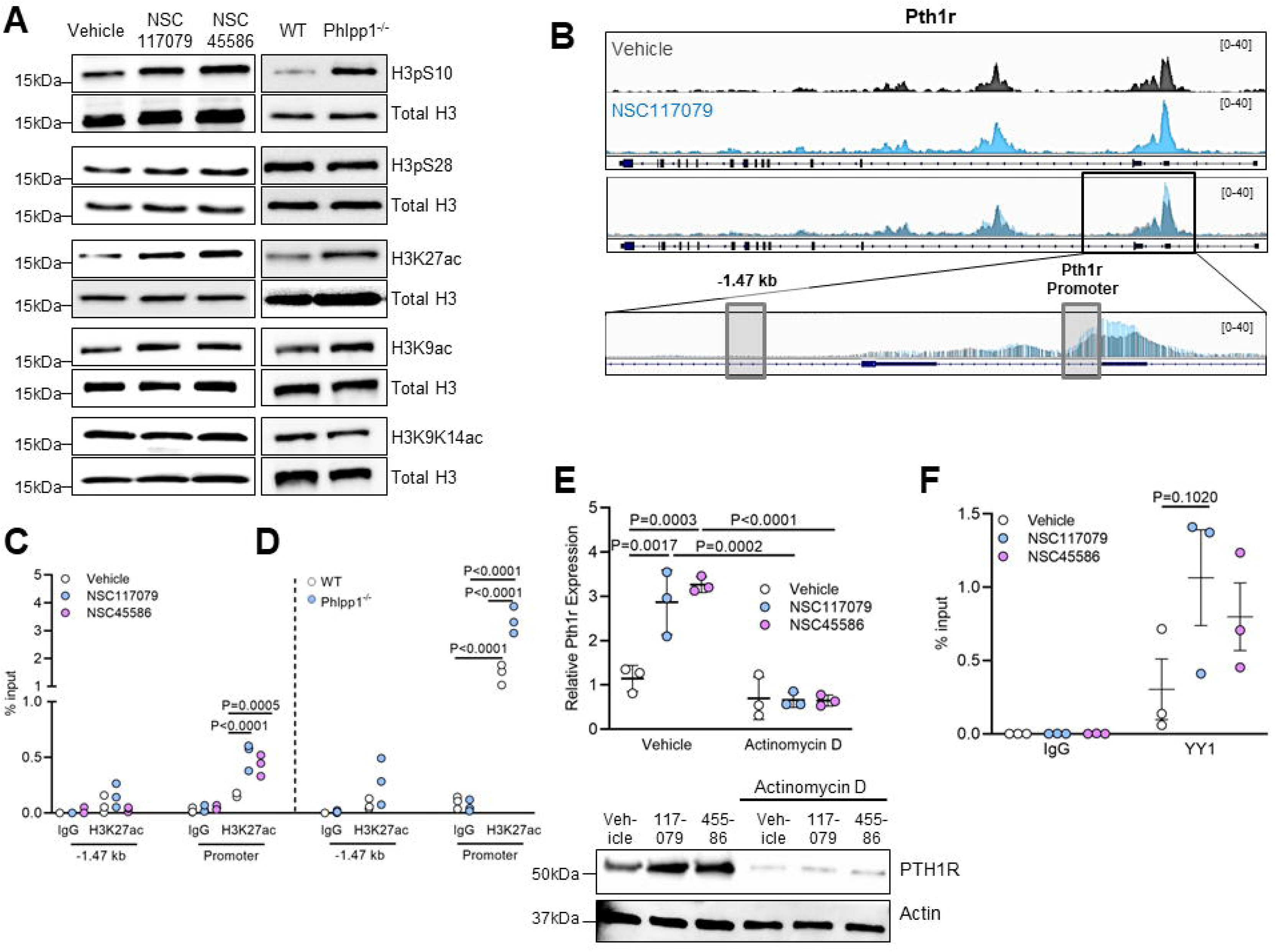
Phlpp1 regulates Pth1r expression through transcription. (A) Primary WT chondrocytes were treated with 25 μM Phlpp inhibitors NSC117079 or NSC45586 for 24 hours. These lysates and those from primary chondrocytes from WT and Phlpp1^−/−^ mice were subjected to Western blotting for phosphorylation of S10 or S28 (pS10 or pS28), as well as K9, K14, and K27 acetylation (ac) of histone 3 (H3). Each blot was also probed with an antibody that recognizes all (total) H3 to control for loading. (B) H3K27ac ChIP-sequencing was performed on WT primary chondrocytes treated for 24 hours with vehicle (DMSO) or 25μm NSC117079. Two sites were identified for analysis by ChIP-qPCR, including in the Pth1r promoter and −1.47kb upstream. (C-D) ChIP-qPCR following pulldown with an H3K27ac antibody was performed on WT primary chondrocytes treated with vehicle or 25μm NSC117079 and NSC45586 after 24 hours (C) or on WT or Phlpp1^−/−^ primary chondrocytes (D). (E) Pth1r mRNA expression and protein levels were measured in primary chondrocytes after incubation with 5 μM transcriptional inhibitor actinomycin D for six hours and subsequent replacement of media containing Phlpp inhibitors for 24 hours. (F) WT primary chondrocytes were treated with vehicle or 25 μm NSC117079 and NSC45586 for 30 minutes. YY1-Pth1r promoter complexes were identified by ChIP-qPCR. Statistically significant differences were determined with two-way ANOVA with Tukey’s post-hoc test.

### PTH(7-34) reverses short limb length in Phlpp1^−/−^ mice

Phlpp1 knockout mice have shorter long bones than WT littermates^(5)^ but similar plasma PTH and urine cAMP concentrations (**Supp Fig 4**). To determine the effect of Pth1r signaling on Phlpp1-mediated long bone growth, whole litters of pups from Phlpp1^+/−^ male and female pairs were injected daily with vehicle or PTH(7-34), which induces receptor internalization but not signaling, from the day of birth through four weeks of age. As expected, Phlpp1^−/−^ mice injected with vehicle had shorter tibiae (**Fig 3A,C,E,G; Supp Fig 5E**) and femurs (**Fig 3B,D,F,H; Supp Fig 5F**), with Phlpp1^+/−^ (HET) showing an intermediate phenotype between WT and Phlpp1^−/−^ mice (**Supp Fig 5A,B,E,F**). Phlpp1^−/−^ mice also had shorter tail-to-snout body length (**Supp Fig 5C,G**; **Supp Fig 6A,C**) and lower body weight (**Supp Figure 5D,H**; **Supp Fig 6B,D**) than WT mice. PTH(7-34) did not alter bone growth in WT mice but rescued the short femur and tibia lengths (**Fig 3**; **Supp Figure 5**) as well as in overall body length and body weight (**Supp Figure 5, 6**) in Phlpp1^−/−^ mice. As males and females showed nearly identical results, male mice were used for all subsequent analyses.

**Figure 3.**
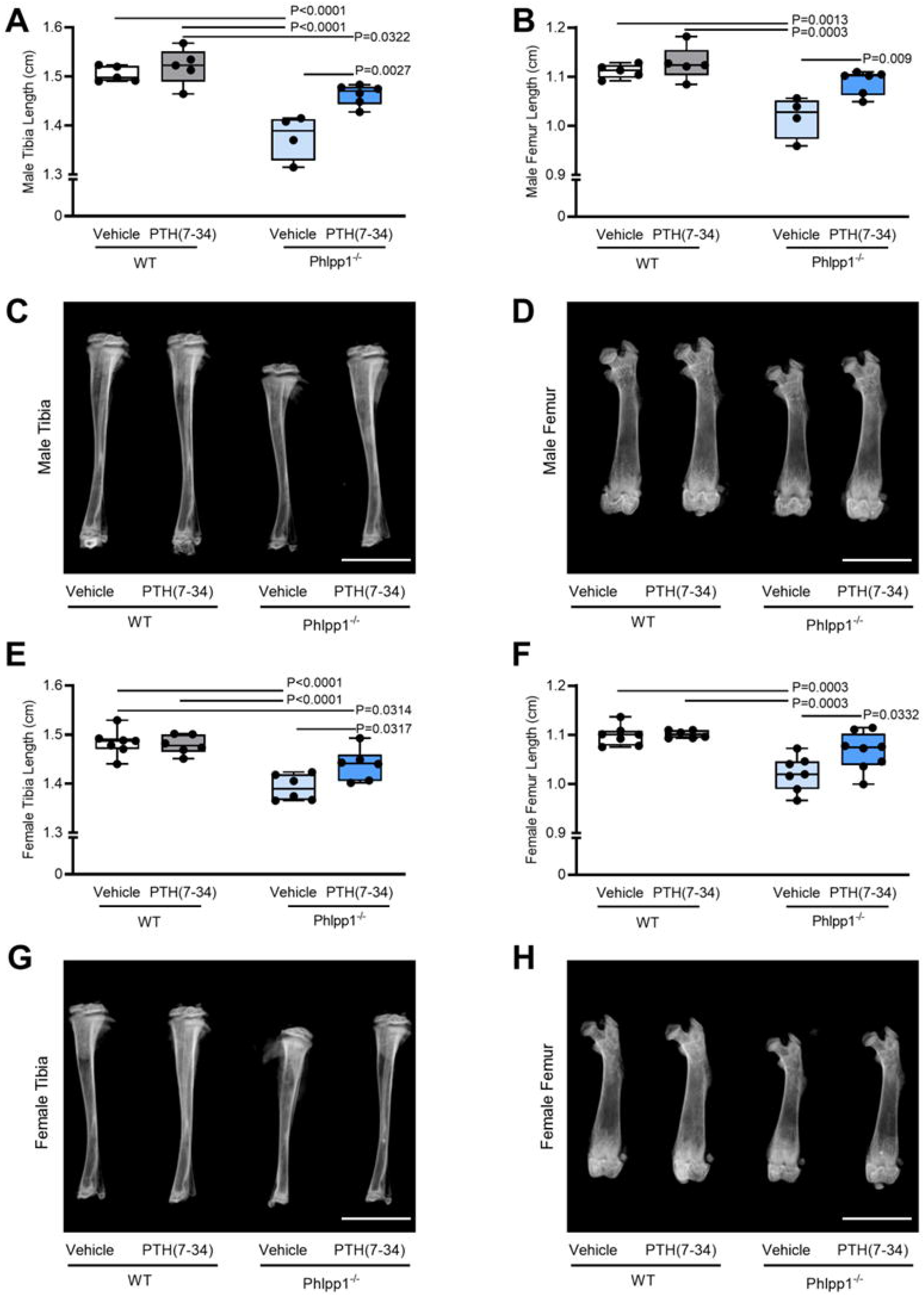
Daily administration of PTH(7-34) reverses short limb length in Phlpp1^−/−^ mice. (A,C) Tibia and (B,D) femur lengths were evaluated in 4-week-old male WT or Phlpp1^−/−^ mice given daily injections of PTH(7-34) (100 mg/kg body weight/day) or vehicle (0.1% BSA in PBS). The same treatments were administered to female WT and Phlpp1^−/−^ mice and tibia (E,G) and femur (F,H) lengths were evaluated at 4 weeks of age. Scale bar = 5 mm. Statistically significant differences were determined with two-way ANOVA with Tukey’s post-hoc test.

### Pth1r and Phlpp1 coordinate to regulate growth plate size

Chondrocyte hypertrophy in the epiphyseal growth plate determines longitudinal bone growth. Four-week-old Phlpp1^−/−^ mice have an increased cell number in the proliferative zone of the growth plate, suggesting delayed entry into hypertrophy and therefore shorter bone length ^(5)^. 3D rendering of the proximal tibial growth plate^(48)^ confirmed that Phlpp1^−/−^ mice had thicker growth plates than WT littermates (**Fig 4A,B**). The difference in growth plate thickness was due to an increase in the size of the proliferative zone, which was attenuated by the administration of PTH(7-34) to Phlpp1^−/−^ mice (**Fig 4C,D; Supp Fig 2B**). Treatment with PTH(7-34) did not have any effect on the proliferative zone of WT mice. The hypertrophic zone area was not statistically different in any of the groups (**Fig 4C,D**) and there was no difference in the expression of Col10a1 as detected by in situ hybridization (**Fig 4E; Supp Fig 2C**). To more directly assess the effects of Phlpp1 deletion and PTH(7-34) administration on cell proliferation, WT and Phlpp1^−/−^ mice were injected with vehicle or PTH(7-34) at the same doses as the 4-week-old mice from postnatal day 1 (P1) daily through P5. BrdU-positive cells were labeled and quantified in the proximal tibial growth plate. As previously reported, Phlpp1^−/−^ P5 mice had a greater number of BrdU+ cells compared to WT mice. PTH(7-34) administration reversed the effect of Phlpp1 deletion, reducing the number of BrdU+ cells in the proximal tibial growth plate to levels similar to WT mice (**Fig 4F,G; Supp Fig 2D**).

**Figure 4.**
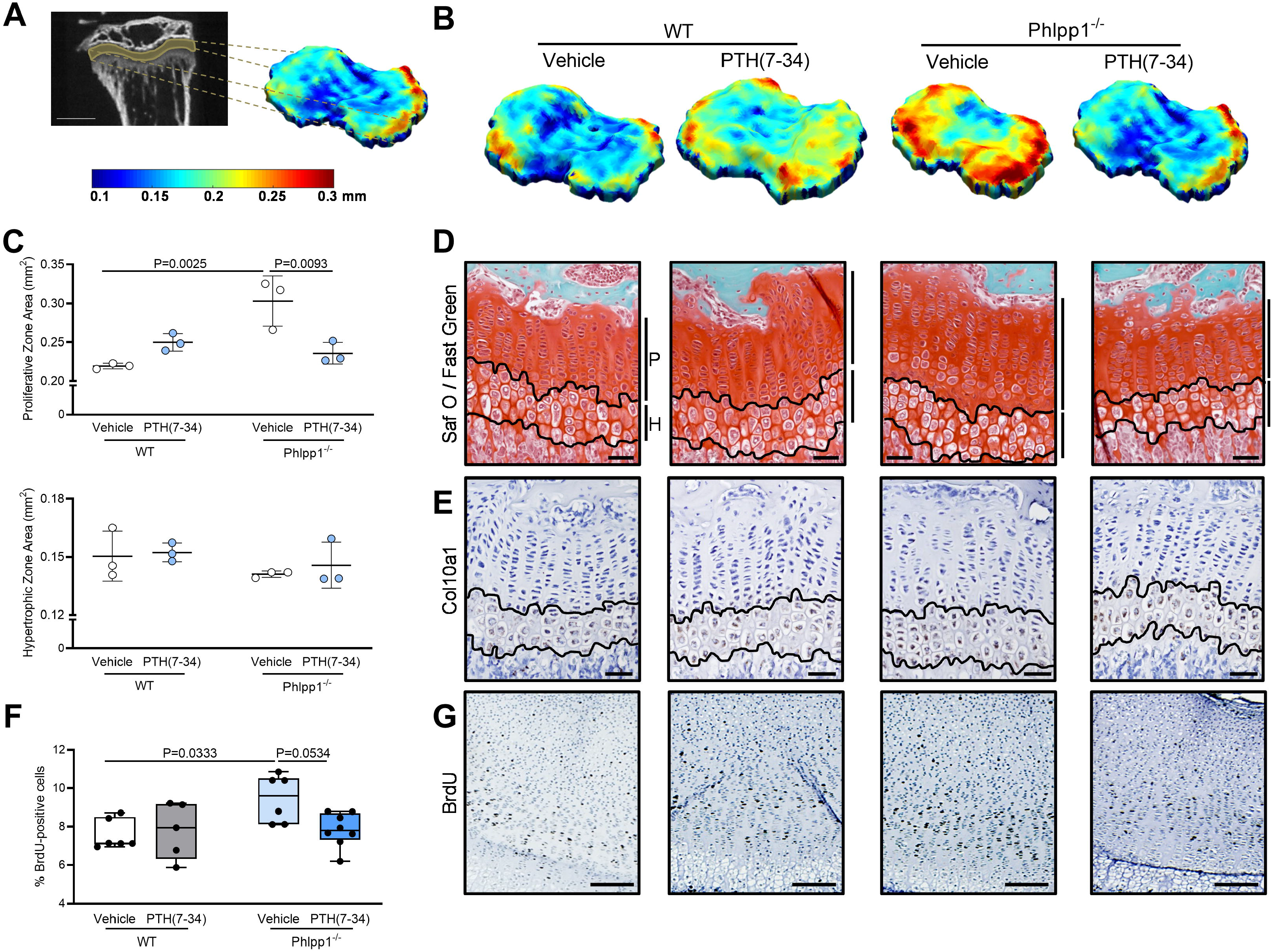
Phlpp1 and Pth1r coordinate to regulate growth plate development. (A,B) 3D renderings of proximal tibial growth plates were generated from microCT scans of 4-week-old WT or Phlpp1^−/−^ male mice given daily injections of PTH(7-34) (100 mg/kg body weight/day) or vehicle. Scale bar on 2D capture = 1mm. Color scale bar is in mm. One representative 3D rendering is shown for each group. (C) Areas of proliferative and hypertrophic zones of the proximal tibia growth plate were quantified. (D) Safranin O / Fast Green staining was performed on the proximal tibial growth plate. Scale bar = 50 μm. Vertical black lines indicate either P = proliferative zone or H = hypertrophic zone. (E) In situ hybridization was performed for Col10a1. Scale bar = 50 μm. (F,G) WT or Phlpp1^−/−^ mice were injected with either vehicle or PTH(7-34) daily as described above from postnatal day 1 (P1) through P5. On P5, BrdU was administered 2 hours prior to euthanasia. (F) Percentage of cells that were BrdU-positive in the proximal tibial growth plate was quantified, as represented in (G). Scale bar in (G) = 50 μm. Statistically significant differences were determined with two-way ANOVA with Tukey’s post-hoc test.

### Daily administration of PTH(7-34) does not affect bone mass

Given the effects of Phlpp1 and Pth1r modulation on bone length, it was pertinent to examine bone mass. Consistent with previous studies^(5)^, Phlpp1^−/−^ mice had lower bone mass than WT littermates. Specifically, Phlpp1^−/−^ mice had lower bone volume / tissue volume (**Fig 5A,B**), fewer trabeculae (**Fig 5C**), and lower trabecular bone mineral density (**Fig 5D**) compared to WT mice. In addition, Phlpp1^−/−^ mice had lower cortical tissue mineral density (**Fig 5E**) and thinner cortical bone (**Fig 5F**). The administration of PTH(7-34) did not affect trabecular bone volume / tissue volume (**Fig 5B**), trabecular number (**Fig 5C**), trabecular bone mineral density (**Fig 5D**), or cortical tissue mineral density (**Fig 5E**) in either WT or Phlpp1^−/−^ mice. However, daily PTH(7-34) administration did reverse the effect of Phlpp1 deletion on cortical thickness (**Fig 5F**). Total tissue cross-sectional area was reduced in Phlpp1^−/−^ mice compared to WT mice, which was reversed by PTH(7-34) administration (**Fig 5G**), as would be expected given the Phlpp1^−/−^ mice have smaller appendicular bones. However, the cortical area fraction was unaffected by Phlpp1 deletion or PTH(7-34) modulation (**Fig 5H**). Trabecular spacing and trabecular thickness were not affected (data not shown).

**Figure 5.**
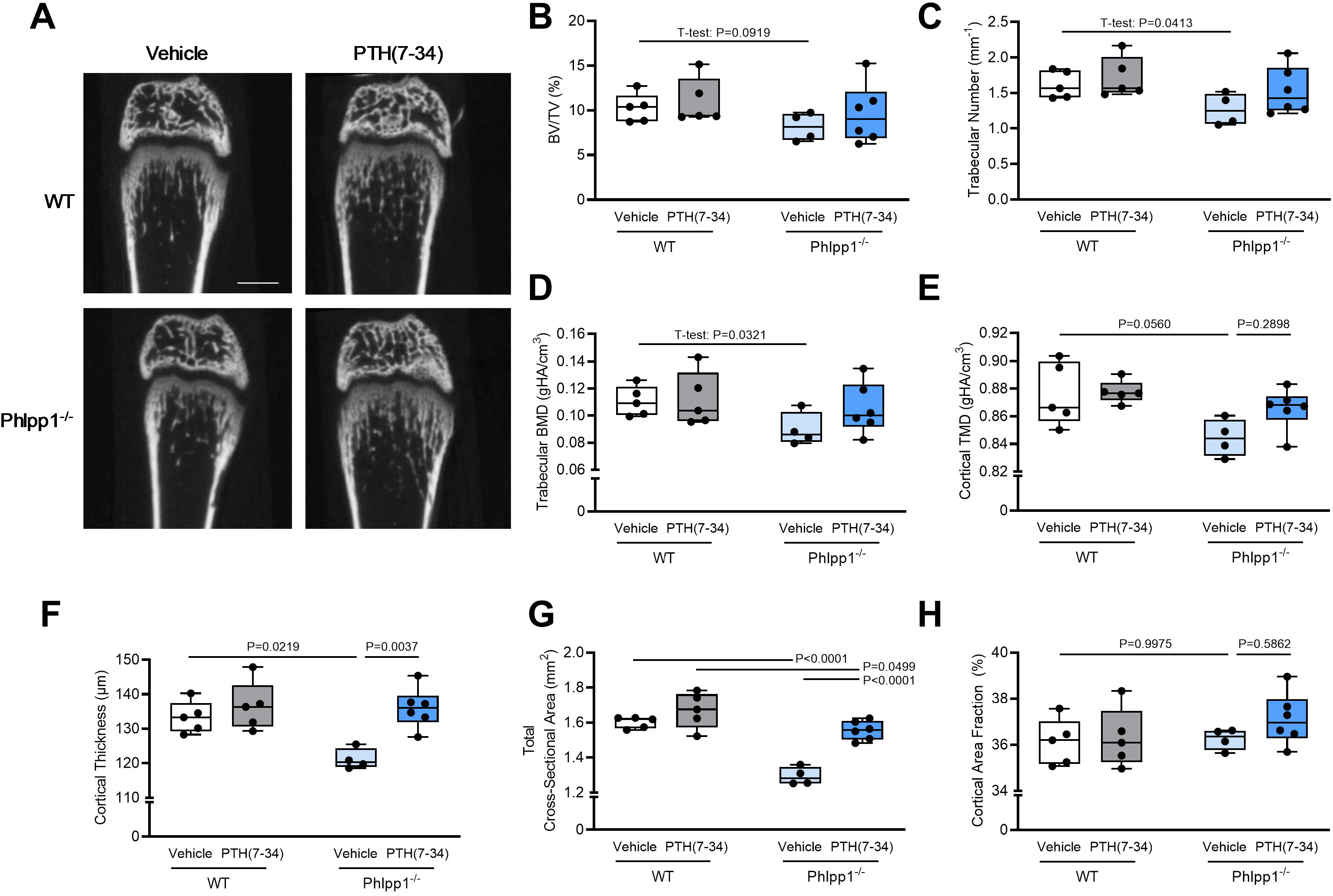
Daily administration of PTH(7-34) does not affect bone mass. (A) 2D reconstructions from microCT of the distal femur of 4-week-old WT or Phlpp1^−/−^ male mice given daily injections of PTH(7-34) (100 mg/kg body weight/day) or vehicle. Scale bar = 1 mm. Trabecular parameters analyzed via microCT were (B) bone volume / tissue volume, (C) trabecular number, and (D) bone mineral density. Cortical parameters included (E) tissue mineral density, (F) cortical thickness, (G) total tissue cross sectional area, and (H) cortical area fraction. Statistically significant differences were determined with two-way ANOVA with Tukey’s post-hoc test or Student’s t test as indicated.

### Phlpp1 inhibition elevates Pth1r signaling cascades

Having established that Phlpp1 regulates Pth1r expression through transcription (**Fig 2**) and controls chondrocyte proliferation in vivo (**Fig 3**), we tested the effects of Phlpp inactivation on known Pth1r signaling targets in vitro. PTH(1-34) was included in these experiment as a positive control. Endogenous PTH and PTH(1-34) bind to Pth1r and stimulate intracellular cAMP, in turn activating PKA and inducing phosphorylation of cAMP-response element-binding protein (CREB) at S133^(52,53)^. IMCs isolated from Phlpp1^−/−^ mice had greater baseline levels of Pth1r and pS133-CREB (**Fig 6A; Supp Fig 1O-Q**). PTH(1-34) enhanced and PTH(7-34) reduced Pth1r and pS133-CREB. Phlpp1^−/−^ IMCs had similar basal cAMP concentrations as WT IMCs, but were more responsive to PTH(1-34) than WT IMCs. PTH(7-34) had no effect (**Fig 6B**). Confluency of Phlpp1^−/−^ chondrocytes increased more rapidly over 48 hours than that of WT monolayer cultures. PTH(1-34) further enhanced confluency of both WT and Phlpp1^−/−^ chondrocyte monolayer cultures, while PTH(7-34) attenuated Phlpp1^−/−^ growth to WT rates (**Fig 6C**). While Phlpp1^−/−^ chondrocytes had only numerically higher MTS activity compared to WT, treatment with PTH(1-34) further enhanced the effects of Phlpp1^−/−^, while PTH(7-34) reversed the effects of Phlpp1^−/−^ on MTS activity (**Fig 6D**). Similar to previous results^(5)^, Phlpp1^−/−^ micromasses have more intense staining of Alcian Blue compared to WT littermates. PTH(1-34) further intensified Alcian Blue staining in Phlpp1^−/−^ chondrocytes. By contrast, treating Phlpp1^−/−^ cells with PTH(7-34) diminished the intensity of Alcian Blue stain to WT levels (**Fig 6E**). Together, these data show that Phlpp1^−/−^ chondrocytes express more Pth1r and are more responsive to PTH ligands.

**Figure 6.**
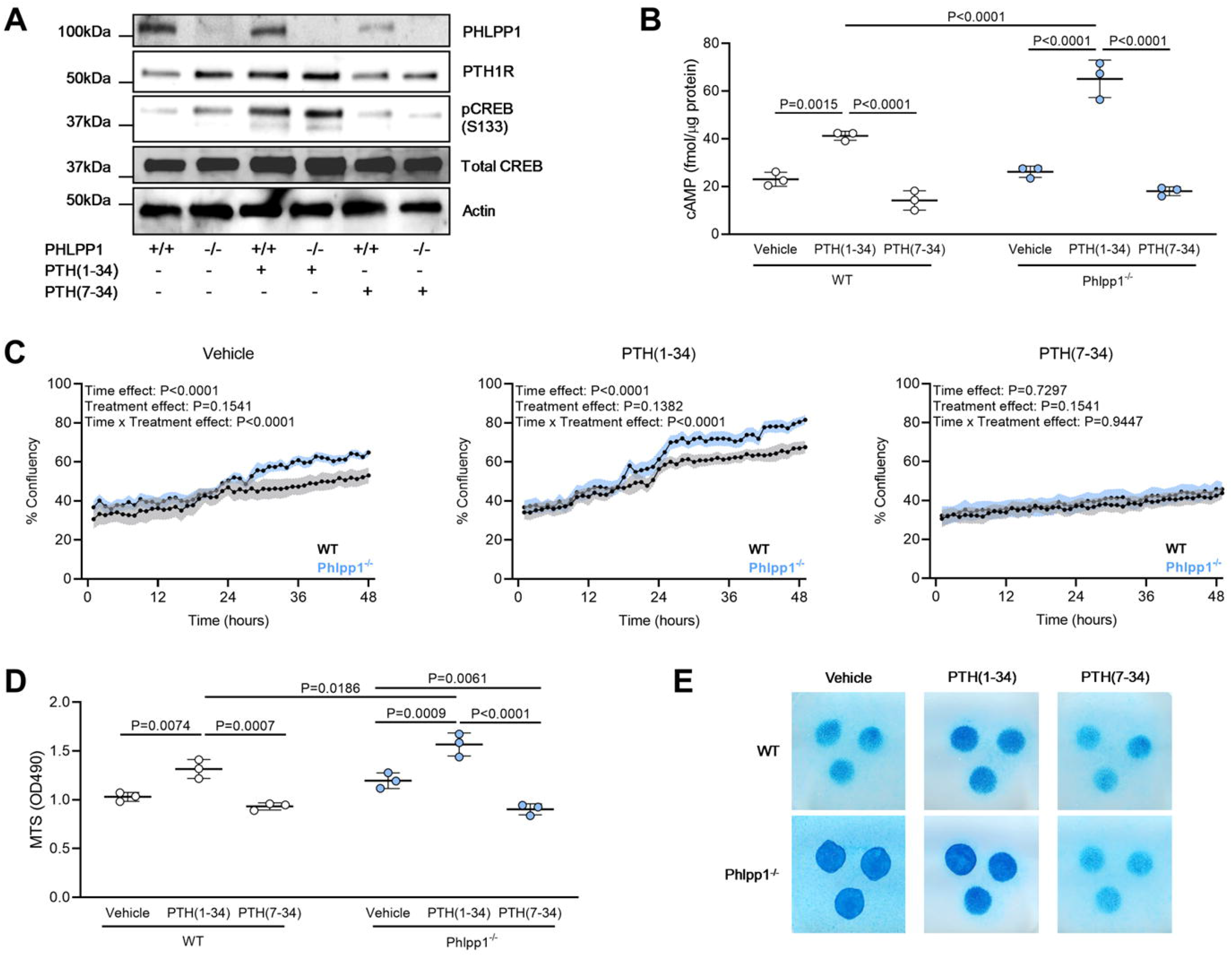
Phlpp1 inhibition elevates Pth1r signaling cascades. (A) Primary chondrocytes from Phlpp1^−/−^ and WT littermates were subjected to Western blotting for Phlpp1, Pth1r, pCREB (S133), total CREB, and actin following 24 hours in culture, with the last 30 minutes in the presence of vehicle (0.1% BSA in PBS), PTH(1-34), or PTH(7-34) (100 nM). (B) cAMP levels in primary chondrocytes from Phlpp1^−/−^ and WT littermates were determined by ELISA following addition of PTH(1-34) or PTH(7-34) (100 nM) for 15 minutes. (C) Confluency of WT and Phlpp1^−/−^ primary chondrocytes was tracked every hour for 48 hours in the absence or presence of PTH(1-34) or PTH(7-34) (10 nM). Data in the three graphs are from the same experiment, but plotted separately for clarity. (D) Primary chondrocytes from Phlpp1^−/−^ and WT littermates were subjected to MTS assay following addition of PTH(1-34) or PTH(7-34) (10 nM) for 24 hours. (E) WT or Phlpp1^−/−^ micromasses were cultured for three days and PTH(1-34) or PTH(7-34) (10 nM) was applied for the next nine days at which time micromasses were stained with Alcian Blue. Statistically significant differences were determined with two-way ANOVA with Tukey’s post-hoc test.

## DISCUSSION

Our results show that greater Pth1r expression and signaling in Phlpp1^−/−^ chondrocytes contributes to growth delays in long bones. Administration of PTH(7-34) for the first four weeks of life reverses the short limb phenotype in the appendicular skeleton of Phlpp1^−/−^ mice. Specifically, PTH(7-34) attenuates the thicker growth plates and large proliferative zones in the epiphyseal growth plate of Phlpp1^−/−^ mice.

Confirming previous results^(5)^, Phlpp1^−/−^ mice have lower bone mineral density than WT mice, and PTH(7-34) has no effect on bone mass or microarchitecture. Regulation of Pth1r by Phlpp1 occurs at the transcriptional level, as H3K27 acetylation is increased at the Pth1r promoter in the Phlpp1-depleted chondrocytes and blocking transcription prevents the increase in Pth1r mRNA and protein levels associated with Phlpp1 inhibition. The transcription factor YY1 may contribute to the increased transcription of Pth1r in Phlpp1 inactivated cells. Additionally, Phlpp1 inhibition increases phosphorylation of Pth1r effector CREB and increases cell proliferation, each of which can be enhanced with Pth1r agonism and attenuated with Pth1r antagonism. Taken together, these results demonstrate that Phlpp1 inhibition increases Pth1r expression and signaling to stunt endochondral ossification.

Pth1r is critical for proper growth plate development. In human populations, both inactivating^(54)^ and activating^(55,56)^ mutations of Pth1r are associated with shortened limbs and abnormal bone mineralization. Mice with a chondrocyte-specific deletion for Pth1r have reduced chondrocyte proliferation, accelerated hypertrophic differentiation, and premature growth plate closure, ultimately resulting in short stature^(35,57)^. Our results show that Pth1r is overexpressed in Phlpp1^−/−^ chondrocytes. In previous studies, Pth1r overexpression in chondrocytes delayed mineralization and decelerated the conversion of chondrocytes from proliferative to hypertrophic states, resulting in shorter limbs^(58)^. Although less dramatic, Phlpp1^−/−^ mice demonstrate a similar phenotype in which bone mineral density is reduced, and the proliferative zone of the growth plate is larger than WT mice.

The altered growth plate phenotype in Phlpp1^−/−^ mice results in shorter femurs and tibiae and this is reversed by administration of PTH(7-34). Our results indicate that Phlpp1-deficiency promotes proliferation through Pth1r signaling and that PTH(7-34) suppression of Pth1r slows proliferation and promotes bone lengthening. Injecting mice with 100 μg/kg body weight/day PTH(7-34) did not fully rescue bone length in every parameter measured. For example, both male and female Phlpp1^−/−^ mice injected with PTH(7-34) had longer tibiae than vehicle-injected Phlpp1^−/−^ mice, but the length was not completely recovered to the control groups. There could be several explanations for these findings. PTH(7-34) was only injected postnatally. PTH signaling is active in utero^(59)^ and it is possible that a basal level of elevated Pth1r signaling had already occurred in Phlpp1^−/−^ mice such that it was only partially attenuated by postnatal administration of PTH(7-34). Other doses and timing schemes of PTH(7-34) administration could be further pursued to determine if there is a more complete rescue. Although our results have focused primarily on chondrocyte dynamics in the growth plate, various hormones, cytokines, and paracrine growth factors work in concert with PTH signaling to determine endochondral ossification and should be further examined in Phlpp1^−/−^ mice^(60)^.

There were no apparent effects of PTH(7-34) on bone mineral density in either WT or Phlpp1^−/−^ mice, with the notable exception of reversing thin cortical bone in Phlpp1^−/−^ mice. PTH signaling plays a significant role in bone mineralization^(61)^ and perturbations in signaling to either enhance or decrease PTH signaling can have profound effects on bone^(62)^. PTH(1-34) increases bone mass when administered intermittently during development and in more mature models^(37,45,63–65)^. Teriparatide (PTH(1-34)) and abaloparatide (PTHrP(1-34)) are widely utilized osteoanabolic treatments that rely on Pth1r signaling for bone accrual^(65)^. Although Phlpp1^−/−^ mice have increased Pth1r expression and signaling, they have lower bone mineral density^(5)^. This is perhaps unsurprising, as constitutively overexpressing Pth1r delays bone mineralization during development^(58)^. Phlpp1 deletion specifically in osteoclasts increases bone mineralization^(7)^ while germline deletion of Phlpp1 decreases bone mass^(5)^, demonstrating that Phlpp1 effects are distinct in different musculoskeletal cell types. The fact that PTH(7-34) was unable to rescue low bone mineralization in Phlpp1^−/−^ mice suggests that Phlpp1 differentially regulates Pth1r signaling in bone cells (osteoblasts, osteoclasts, and osteocytes) compared to chondrocytes. Future studies should probe the relationship between Phlpp1 and Pth1r in individual musculoskeletal types to elucidate this relationship beyond chondrocytes.

In vitro, our results recapitulate the in vivo relationship between Phlpp1 inhibition and enhanced Pth1r signaling in chondrocytes. While agonism of Pth1r with PTH(1-34) enhances Phlpp1^−/−^ responses, attenuation of PTH signaling with PTH(7-34) can reverse the effects of Phlpp inhibition. Phlpp1-depleted cells show increased phosphorylation of CREB and treating Phlpp1^−/−^ cells with PTH(1-34) further enhanced pCREB, while PTH(7-34) reversed this indicator of Pth1r signaling. PTH(1-34) is more responsive in Phlpp1^−/−^ cells that have higher concentrations of the Pth1r receptor available for PTH(1-34) binding, as cell metabolic activity and cAMP concentrations were highest in Phlpp1^−/−^ cells treated with PTH(1-34). Similarly, cell proliferation, as well as cartilage matrix deposition, were increased in Phlpp1^−/−^ cells compared to wild type littermates, with PTH(7-34) attenuating elevated proliferation and chondrogenesis associated with Phlpp1 inhibition. In vivo, BrdU incorporation in the growth plate of P5 mice confirmed the in vitro findings. Further enhancement of cell proliferation as a result of Phlpp1 inhibition has potential implications beyond chondrocytes. Phlpp1 deletion promotes cell proliferation in a model of intervertebral disc degeneration^(19)^ and regulation of cell apoptosis by Phlpp1 is implicated in ischemic brain and spinal cord injury^(20,22)^, colitis^(21)^, cardiac dysfunction^(66)^, pancreatic beta-cell survival^(67)^, and tumor activity^(8)^. As such, enhancement of signals associated with Phlpp1 inhibition could have a profound impact on various physiological and disease processes.

Phlpp1 modulates Pth1r signaling through chromatin remodeling of the ubiquitous Pth1r promoter resulting in increased transcription, as well as greater activation of Pth1r signaling^(68)^. The mechanisms responsible for increased transcription of Pth1r via Phlpp inhibition remain to be determined, but appear to involve a modest increase in existing transcriptional events. A variety of transcription factors are known to bind to the Pth1r promoter and may be active in Phlpp-inhibited chondrocytes^(68–72)^. Numerous potential transcription factor binding sites within the 2,000 base pair H3K27ac enrichment peak in the Pth1r promoter were identified and are summarized in **Supplementary Table 1**^(73)^. One promising transcription factor, YY1, showed increased binding at the Pth1r promoter in response to Phlpp inhibition. Ying Yang 1 (YY1) is a dual function transcription factor that regulates both transcriptional activation and repression and has been implicated in a host of cellular processes, including differentiation, DNA repair, cell division, and cell survival^(74)^. Additionally, YY1 facilitates the interaction of active enhancers and promoter-proximal elements, serving as a general feature of mammalian gene control^(75)^. In chondrocytes, Phlpp inhibitors increase binding of YY1 to the Pth1r promoter, suggesting that YY1 acts as a transcriptional activator in Phlpp-inactivated chondrocytes. However, because a large number of factors could associate with this region of the Pth1r promoter, it is likely that Phlpp1 repression of H3K27ac influences many additional transcription factors and co-regulators.

The relationship between Phlpp1 and Pth1r has applications beyond endochondral ossification during development. Pth1r^−/−^ mice have spontaneous cartilage degeneration and develop more severe post-traumatic osteoarthritis in the knee than wild type littermates, with greater chondrocyte hypertrophy^(76)^. By contrast, Phlpp1^−/−^ mice and WT mice given an intra-articular injection of NSC117079 into the knee joint are protected from the development of post-traumatic osteoarthritis, with improved allodynia and functional impairments. As such, Phlpp inhibitors have potential as novel disease modifying osteoarthritis drugs (DMOADs). Human recombinant PTH(1-34) (teriparatide) is currently in clinical trials as a DMOAD, following preclinical results of reduced cartilage degeneration and induction of matrix regeneration^(77)^. As our findings indicate a synergistic relationship between Phlpp1 inhibition and enhanced Pth1r signaling, it is possible that combinations of Phlpp1 inhibitors and PTH(1-34) could be effective promoters of cartilage regeneration.

In summary, our results demonstrate a novel molecular relationship in which Phlpp1 inhibition enhances Pth1r expression and signaling in chondrocytes. Repression of Pth1r signaling reversed both the stunted growth of the appendicular skeleton characteristic of Phlpp1^−/−^ mice, and the increased cell proliferation of Phlpp1^−/−^ chondrocytes, both in vitro and in the proliferative zone of the epiphyseal growth plate. Our findings have implications on multiple processes, including endochondral ossification during development and musculoskeletal diseases such as osteoarthritis. Phlpp1 and Pth1r signaling is not restricted to the bone environment, as PTH controls mineral homeostasis^(78)^ among other processes, and Phlpp1 is expressed throughout the mammalian body^(6)^. As such, the Phlpp1-Pth1r relationship could have a significant impact in the field of musculoskeletal health and beyond.

## ACKNOWLEDGEMENTS

The authors thank Xiaodong Li and Carys Turner for technical assistance. The Mayo Clinic X-Ray Imaging Resources Core Facilities were essential for collection of the microCT data. We are grateful to the Mayo Clinic Medical Genome Facility Sequencing Core for performing and helping with the analysis of ChIP-Sequencing. This work was supported by research and training grants from the National Institutes of Health (AR068103, AR056950, DK07352, AR065397, AR072634).

Authors’ roles: Study design: EWB, JJW. Study conducted: all authors. Data collection and analysis: all authors. Data interpretation: SRW, ELT, EWB, JJW. Writing – original draft: SRW. Revising and approving final version of manuscript: all authors. Supervision: EWB, JJW. Project leadership: SRW, ELT, EWB, JJW. Funding acquisition: EWB, JJW. SRW and JJW take responsibility for the integrity of the data analysis.

## FIGURE LEGENDS

**Supplementary Figure 1.**
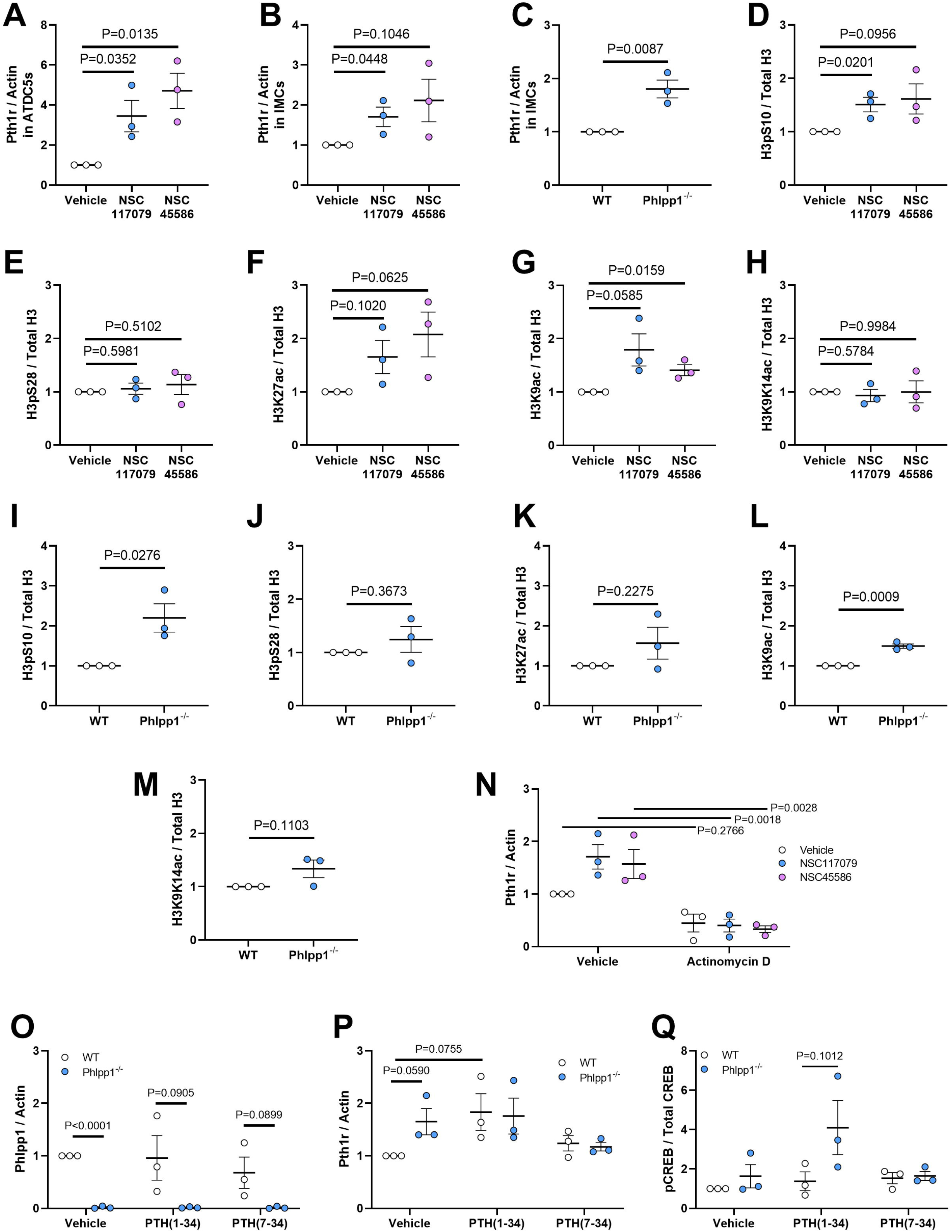
Quantified Western Blots. (A-C) Pth1r protein levels were measured in (A) ATDC5s and (B) WT primary chondrocytes after incubation with vehicle or Phlpp inhibitors (25μm NSC117079 or NSC45586) for 24 hours, as well as in (C) WT or Phlpp1^−/−^ primary chondrocytes. Actin was the loading control. (D-M) (D-H) WT primary chondrocytes were treated with Phlpp inhibitors for 24 hours. (I-M) Primary chondrocytes from WT and Phlpp1^−/−^ were grown in culture for 24 hours. The protein levels measured were (D,I) H3pS10, (E,J) H3pS28, (F,K) H3K27ac, (G,L) H3K9ac, and (H,M) H3K9K14ac. Total histone 3 (H3) was the loading control. (N) Pth1r protein levels were measured in primary chondrocytes after incubation with 5 μM actinomycin D for six hours and subsequent replacement of media containing Phlpp inhibitors for 24 hours. (O-Q) Primary chondrocytes from Phlpp1^−/−^ and WT mice were subjected to Western blotting for Phlpp1, Pth1r, and pCREB (S133) with actin and total CREB as the loading controls following 24 hours in culture, with the last 30 minutes in the presence of vehicle, PTH(1-34) (100 mM), or PTH(7-34) (100 nM).

**Supplementary Figure 2.**
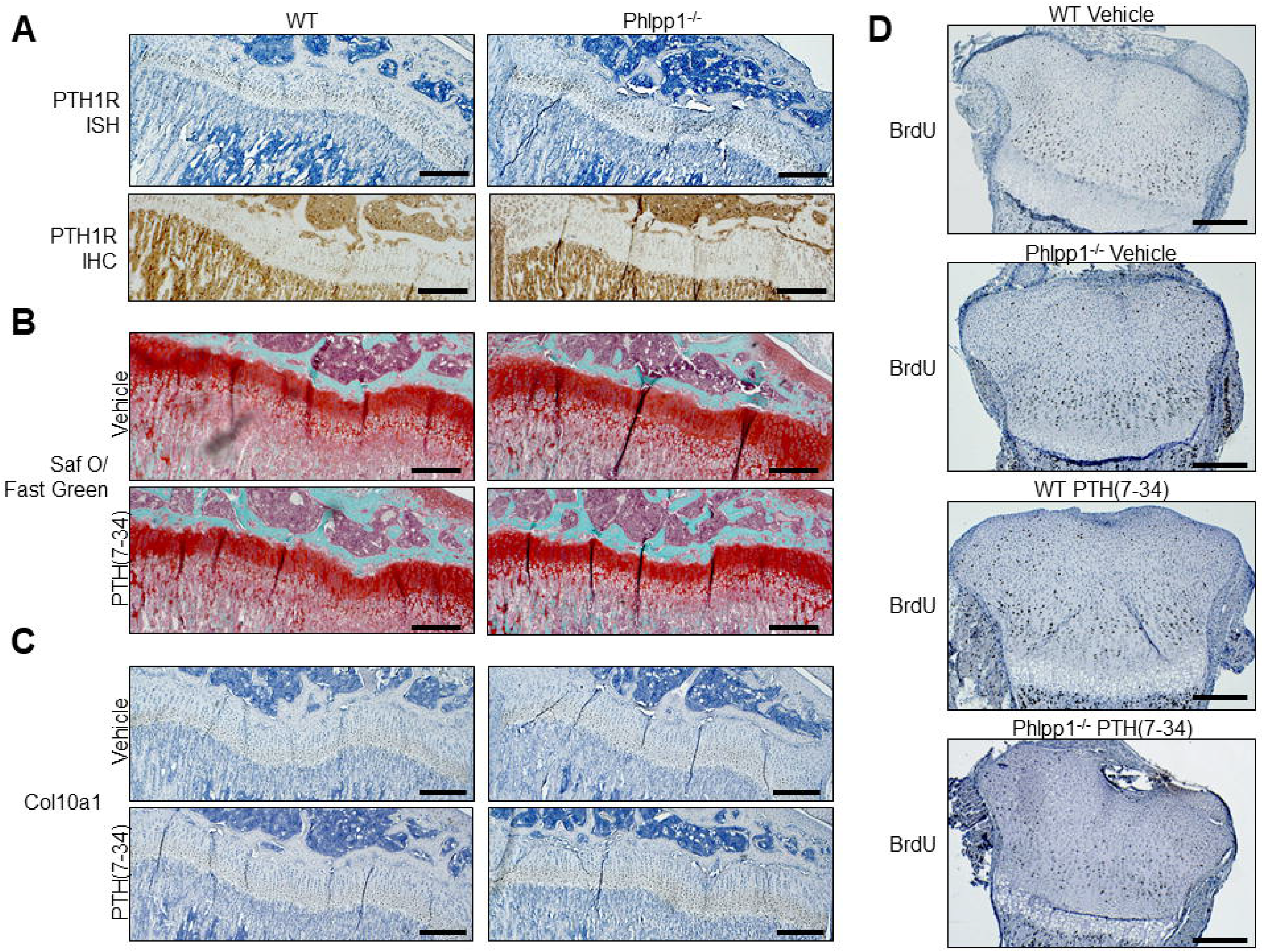
Low magnification images of the growth plate. 5x magnification images of (A) the growth plates of 4-week-old WT or Phlpp1^−/−^ male mice after performing in situ hybridization (ISH) or immunohistochemistry (IHC) to detect Pth1r. Scale bar = 100 μm. (B,C) Four-week-old WT or Phlpp1^−/−^ mice were given daily injections of PTH(7-34) (100 mg/kg body weight/day) or vehicle. The proximal tibial growth plate was (B) stained with Safranin O / Fast Green and (C) probed for Col10a1 using ISH. (D) WT or Phlpp1^−/−^ mice were injected with either vehicle or PTH(7-34) daily as described above from postnatal day 1 (P1) through P5. On P5, BrdU labeling reagent was administered 2 hours prior to euthanasia and BrdU-positive cells were identified.

**Supplementary Figure 3.**
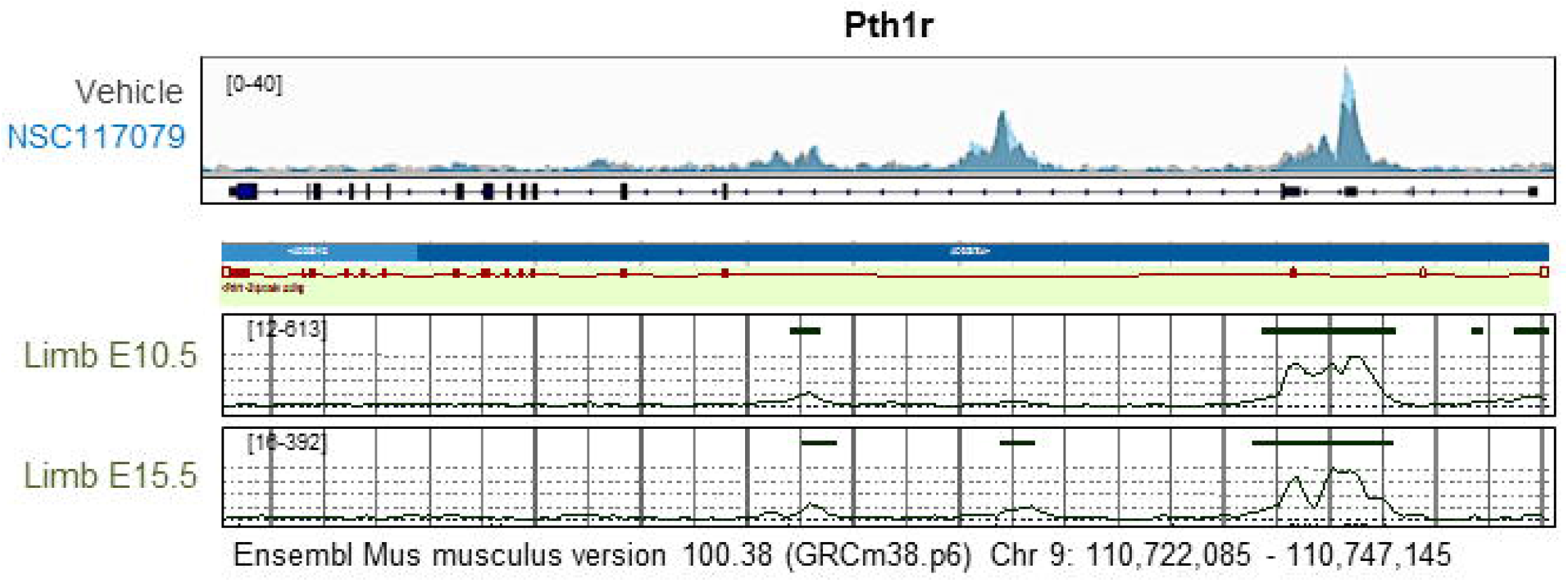
Small-molecule Phlpp inhibitors increase H3K27ac in Pth1r. H3K27ac ChIP-sequencing was performed on WT primary chondrocytes incubated for 24 hours with vehicle (DMSO) or 25μm NSC117079. Data were aligned with publicly available ChIP-Seq datasets from E10.5 and E15.5 limbs for Chr 9: 100,722,085 – 110,747,145. Limb data was retrieved from Ensembl for Mus musculus, version 100.38 (GRCm38.p6).

**Supplementary Figure 4.**
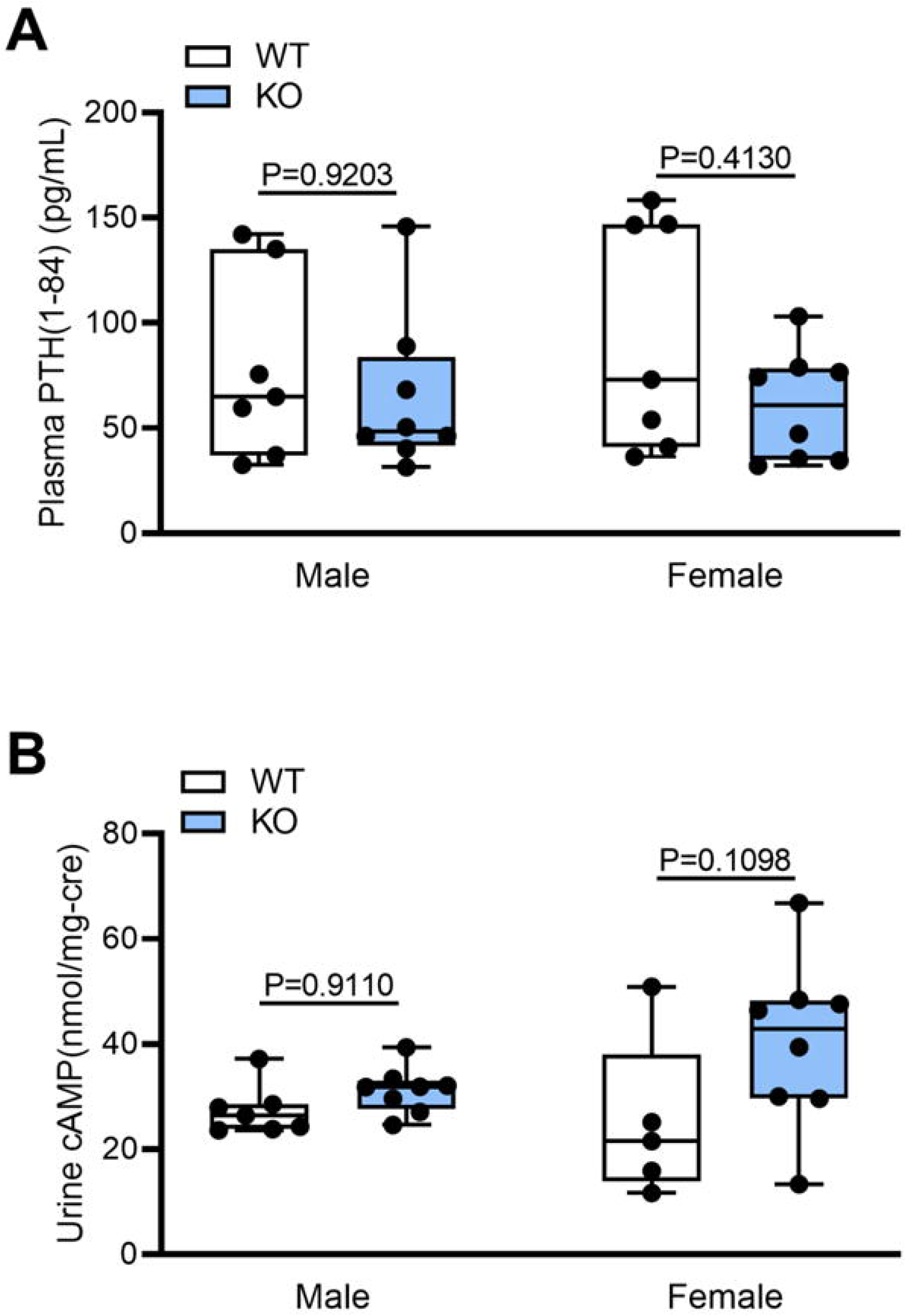
Plasma PTH and urine cAMP are the same in WT and Phlpp1^−/−^ mice. (A) Plasma intact PTH(1-84) and (B) urine cAMP corrected for urine creatinine were measured in samples taken from four-week-old WT and Phlpp1^−/−^ mice. Statistically significant differences were determined via one-way ANOVA with Tukey’s post-hoc test.

**Supplementary Figure 5.**
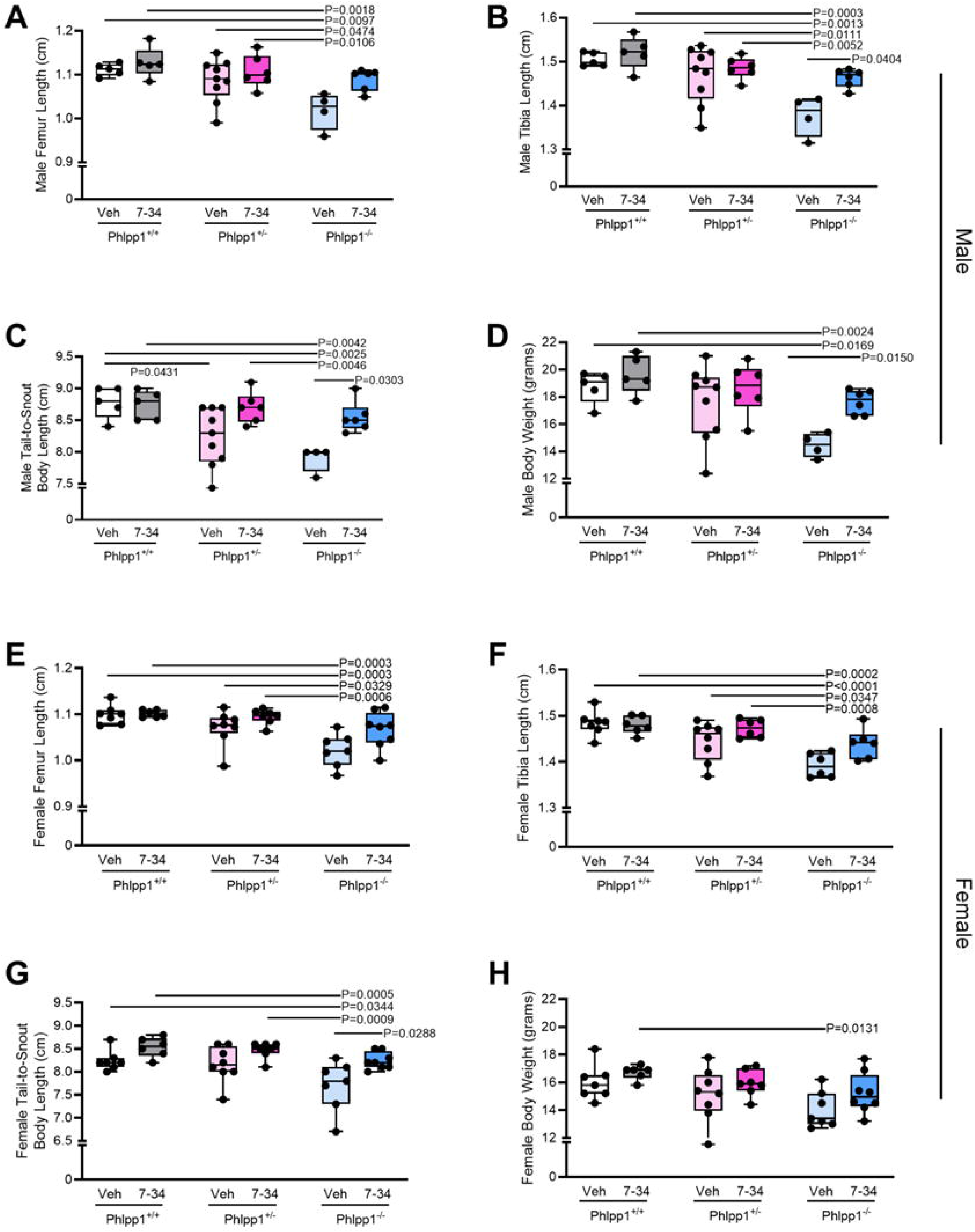
Daily administration of PTH(7-34) has an intermediate effect on rescuing limb length in Phlpp1 heterozygotes compared to WT or Phlpp1^−/−^ mice. (A,E) Femur length, (B,F) tibia length, (C,G) body length, and (D,H) body weight were evaluated in 4-week-old male and female WT (Phlpp1^+/+^), HET (Phlpp1^+/−^), or KO (Phlpp1^−/−^) mice given daily injections of PTH(7-34) (100 mg/kg body weight/day) or vehicle (0.1% BSA in PBS). Statistically significant differences were determined with two-way ANOVA with Tukey’s post-hoc test.

**Supplementary Figure 6.**
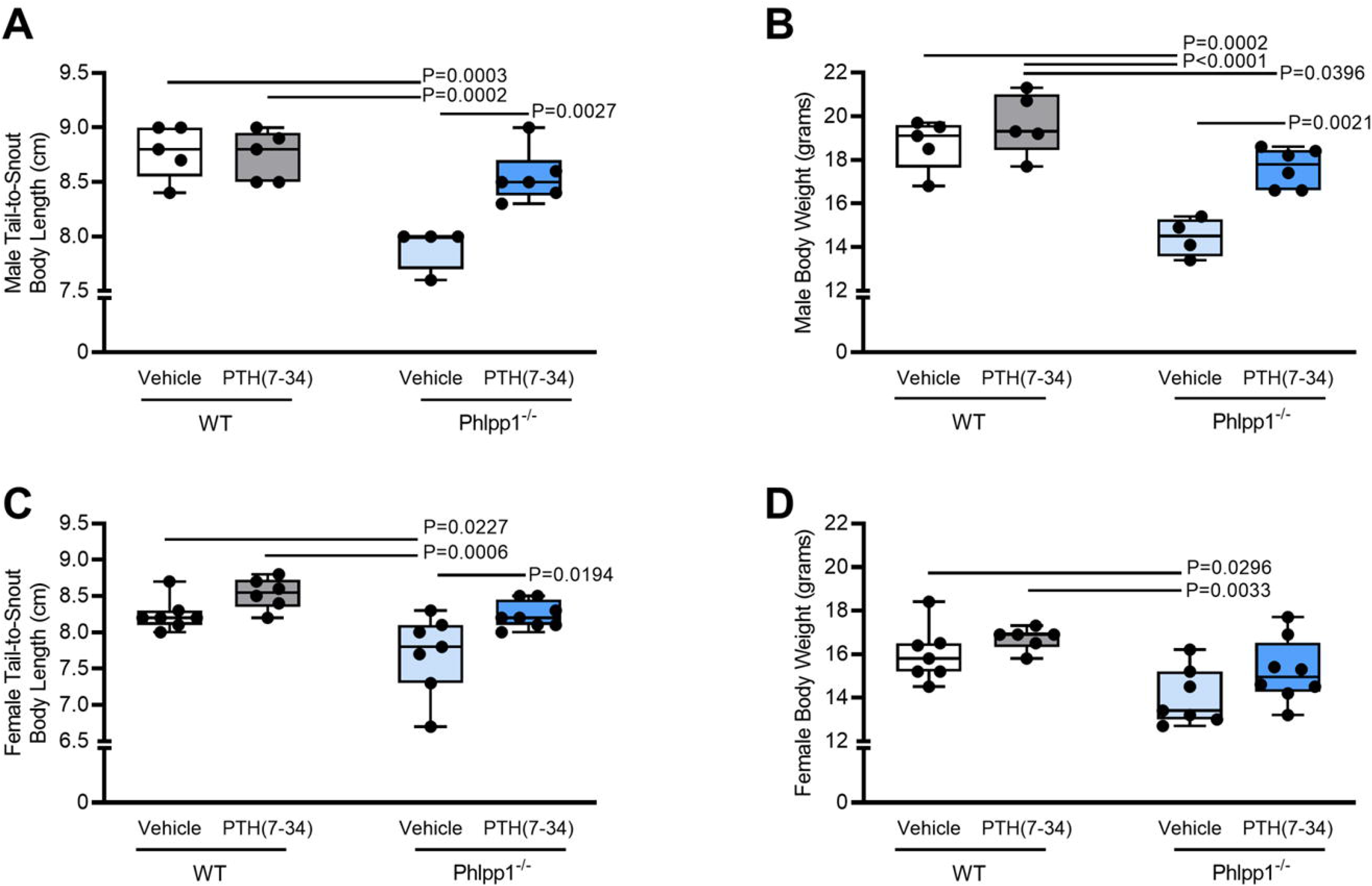
Daily administration of PTH(7-34) reverses short body length and low body weight in Phlpp1^−/−^ mice. (A,C) Tail-to-snout body length and (B,D) body weight were evaluated in 4-week-old male and female WT or Phlpp1^−/−^ mice given daily injections of PTH(7-34) (100 mg/kg body weight/day) or vehicle (0.1% BSA in PBS). Statistically significant differences were determined with two-way ANOVA with Tukey’s post-hoc test.

## TABLES

**Supplementary Table 1.**
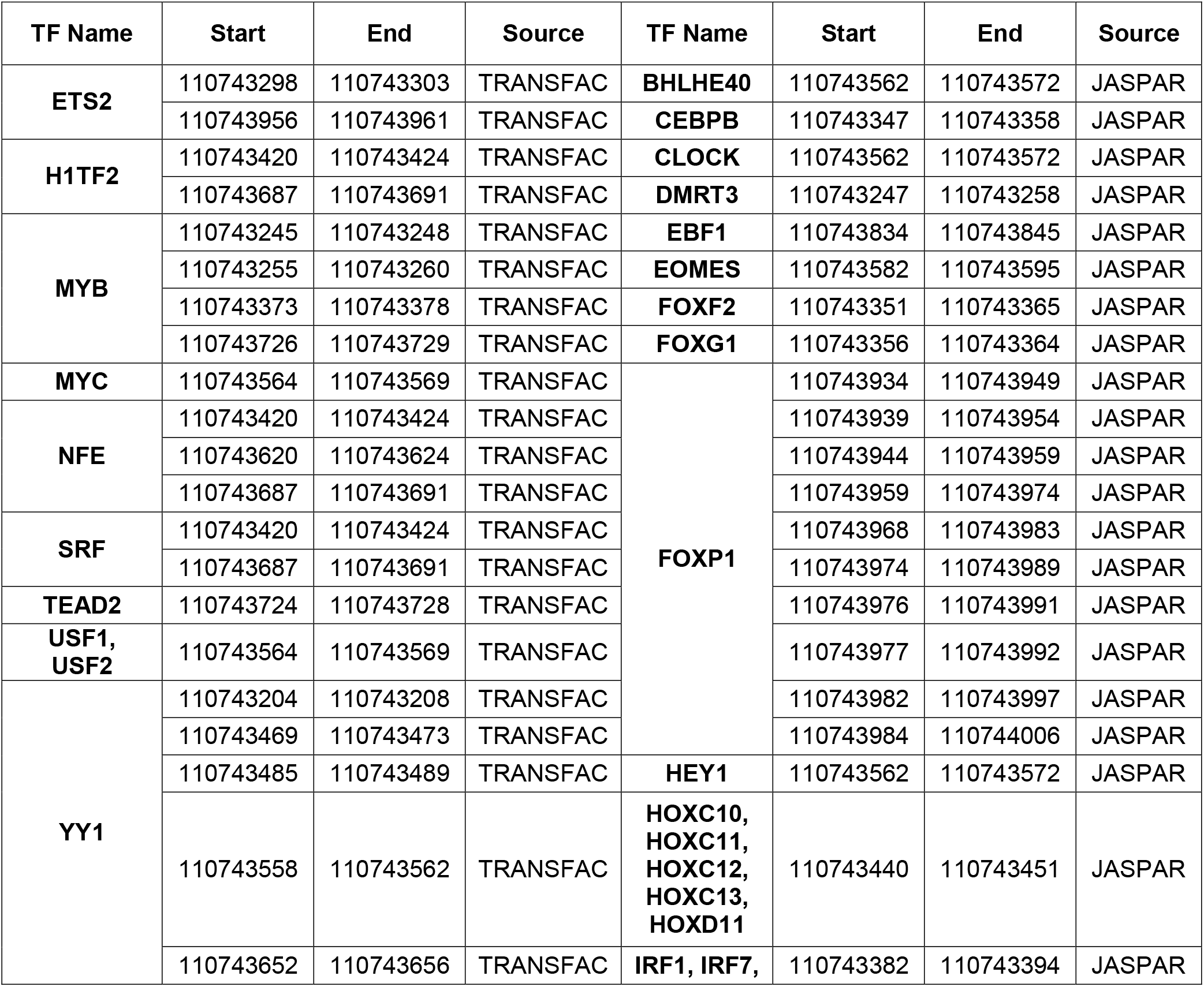
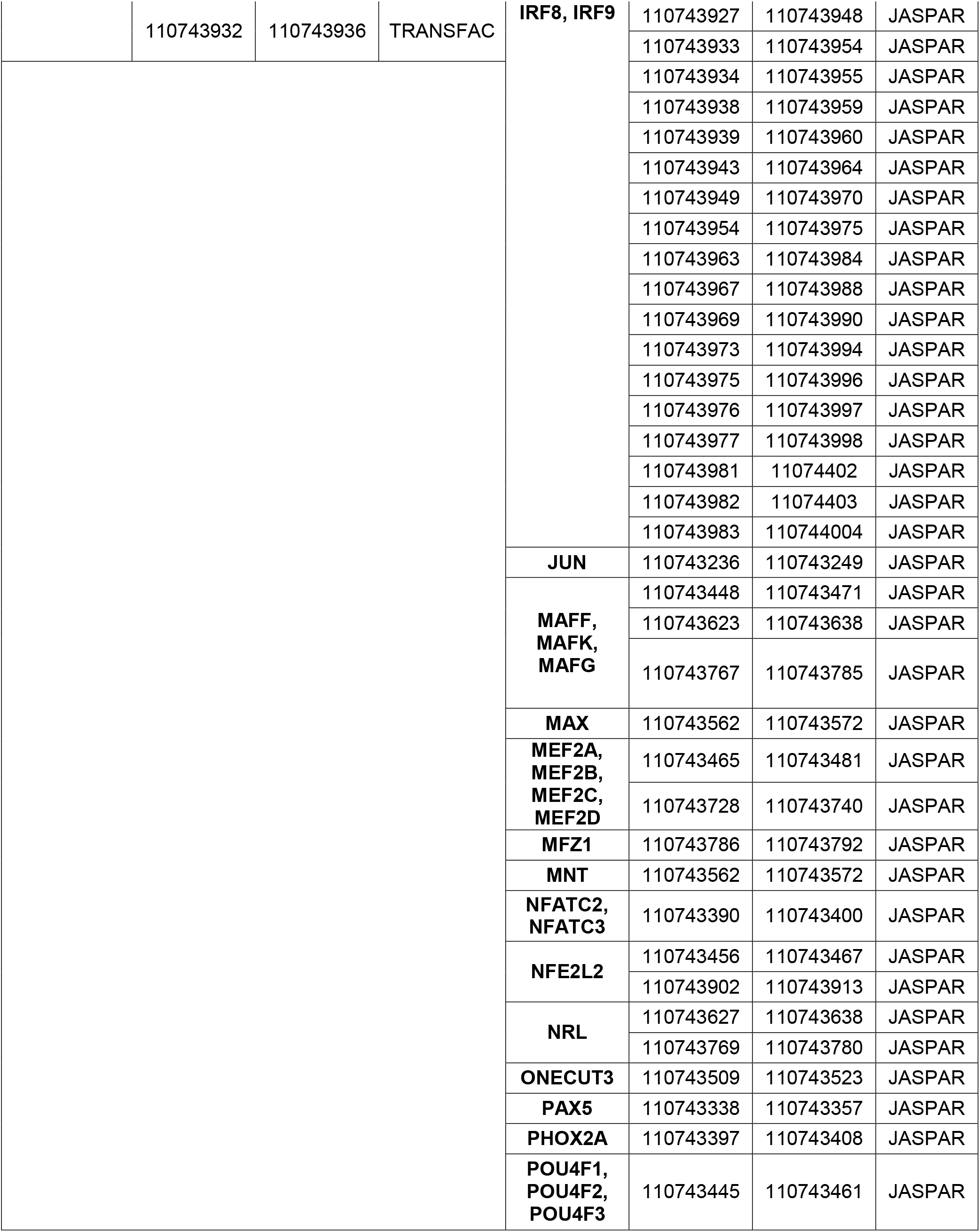
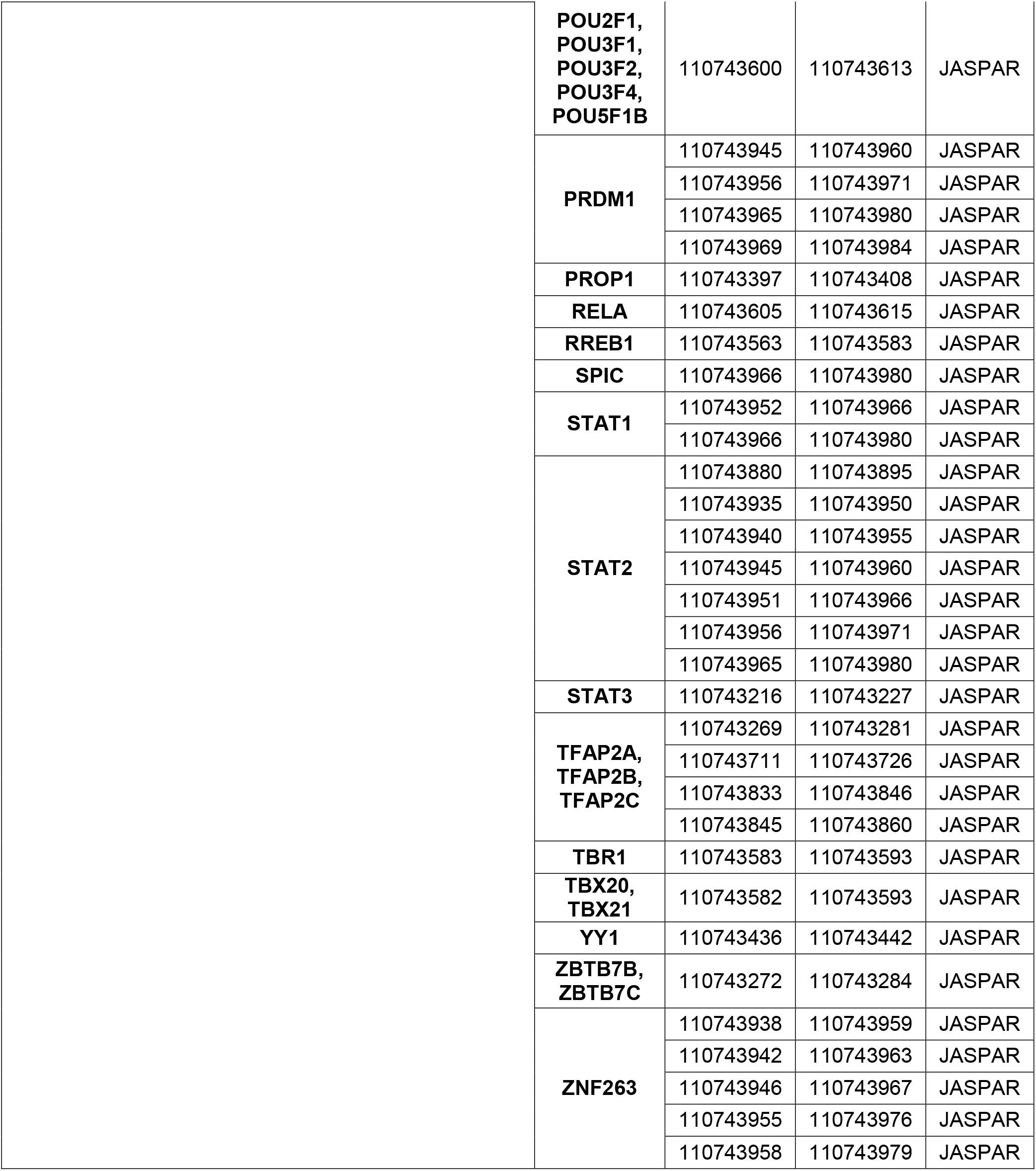
Candidate transcription factor binding sites within the Pth1r promoter. Primary chondrocytes were treated with NSC117079 for 24 hours and analyzed by ChIP-Seq. The DNA sequence under the H3K27Ac peak in the Pth1r promoter (chromosome 9: 110742000-110744000) was used to search EnhancerDB (http://lcbb.swjtu.edu.cn/EnhancerDB/) and identify candidate transcription factors binding to this region.

**Supplementary Table 2.**
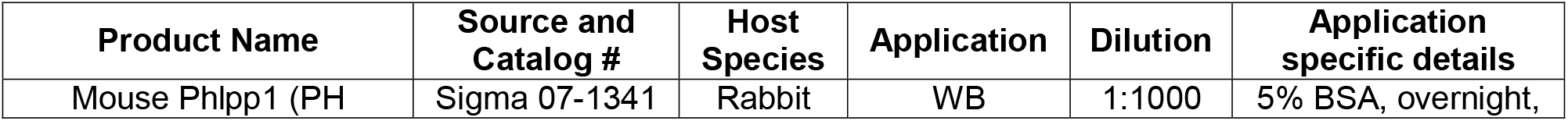
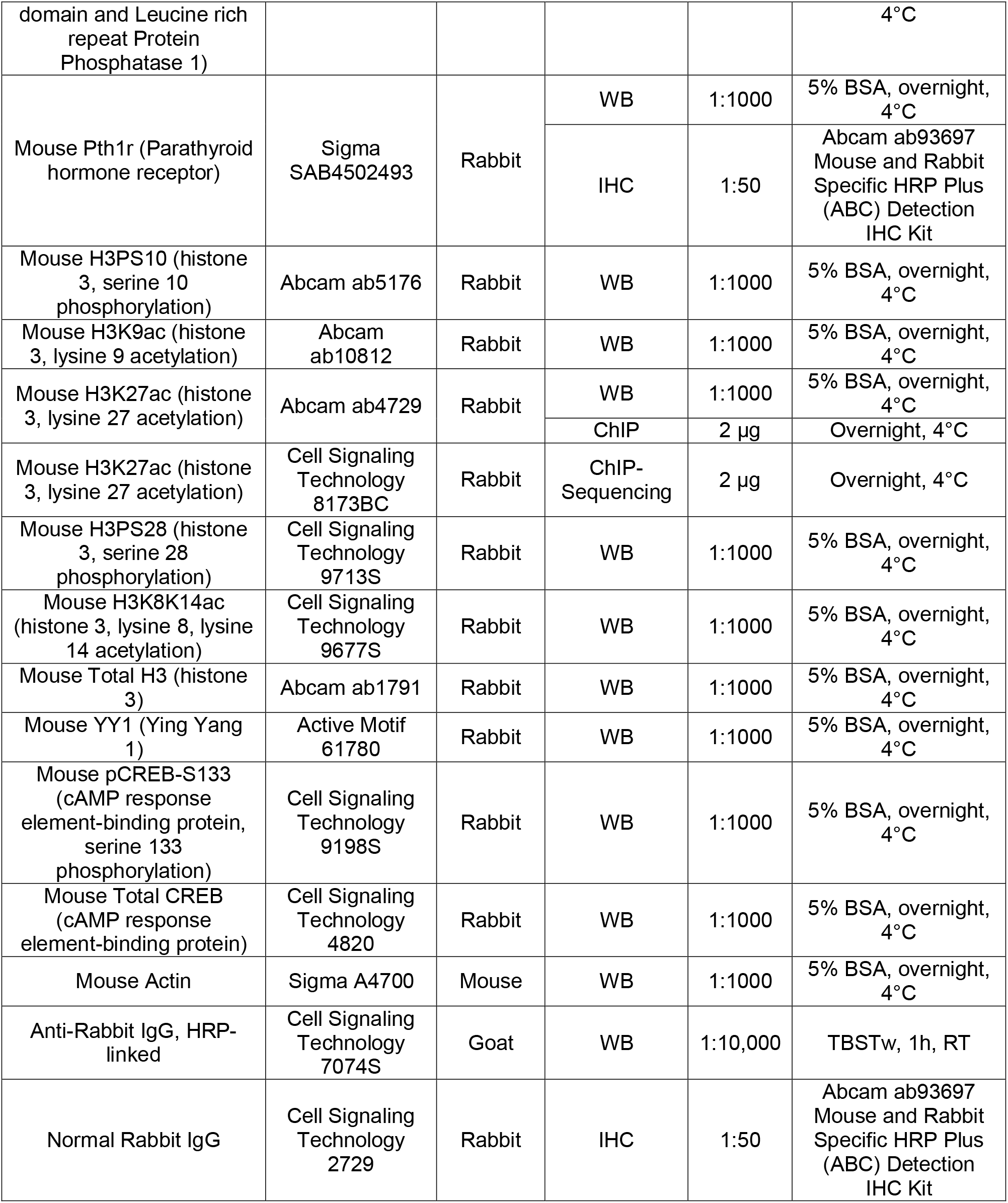
Antibodies.

**Supplementary Table 3.**
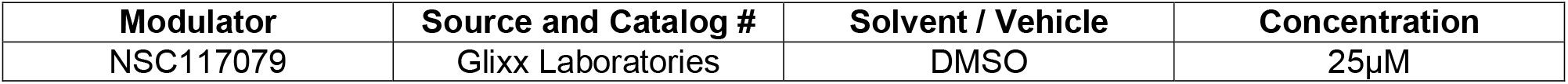
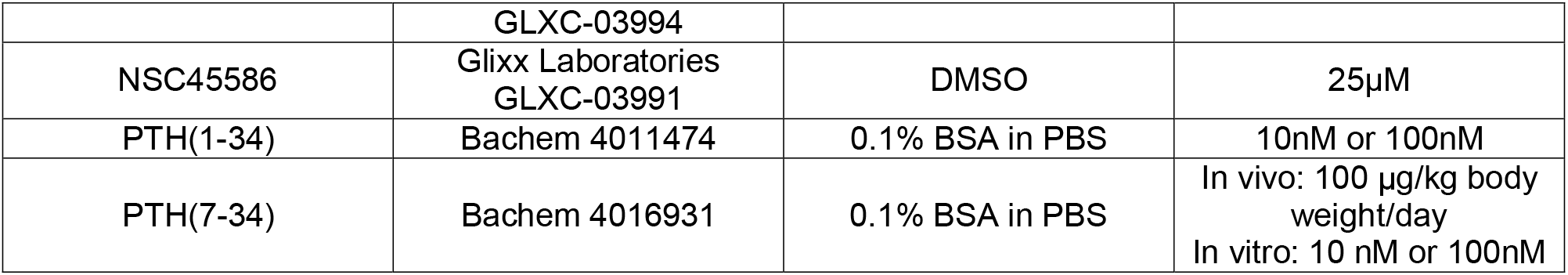
Biological Modulators.

